# Soybean-SCN duel: Novel insight into Soybean’s Resistant Responses to *Heterodera glycines*

**DOI:** 10.1101/2023.05.22.541756

**Authors:** Sepideh Torabi, Soren Seifi, Jennifer Geddes-McAlister, Albert Tenuta, Owen Wally, Davoud Torkamaneh, Milad Eskandari

## Abstract

Soybean cyst nematodes (SCN, *Heterodera glycines* Ichinohe) are a significant threat to soybean production globally, causing severe yield losses and necessitating the development of effective strategies to combat this devastating nematode disease. This study presents dual RNA-seq analyses of the three most common SCN-resistant lines (Plant Introduction (PI) 437654, 548402, and 88788) and the susceptible line Lee74 against the SCN HG type 1.2.5.7 to identify the mechanisms of resistance and virulence genes involved in resistance breakdown. Transcriptomic and pathway analyses reveal the activation of the phenylpropanoid pathway, MAPK signaling pathway, plant hormone signal transduction, and secondary metabolite pathways in the resistance mechanisms. PI 437654, which exhibited robust resistance (female index, FI=0%), demonstrated unique gene expression associated with cell wall reinforcement, oxidative enzymes, ROS scavengers, and Ca+2 sensors governing the salicylic acid (SA) biosynthesis process, indicating its key defense mechanism. Moreover, using different hosts with varying levels of immunity and a susceptible line provided insights into SCN pathogenesis and how *H. glycine* overcomes different layers of host immunity by modulating its virulence genes. This research provides novel insights into the molecular mechanisms underlying soybean-SCN interactions and identifies potential targets for developing strategies to manage this devastating nematode disease.

## Introduction

Nematodes are multicellular animals and belong to the superphylum Ecdysozoa. Nematodes are parasitic and free-living worms that can shed their cuticle to grow (Lambert, K., & Bekal, S., 2002). Plant parasitic nematodes are recognized as major agricultural pathogens (Sasser, J. N. (1987). One of the most destructive plant-parasitic nematodes that can result in significant yield losses and economic damage is soybean cyst nematode (SCN), caused by *Heterodera glycine* (HG) Ichinohe (Carter et al. 2018). SCN poses a serious threat to world soybean production, causing annual yield losses in excess of $1.5 billion in North America (Bradley et al. 2021).

Soybeans, as with other plants, fight-off pathogens through initiation of an array of defence mechanisms. A plant defence system is a complex system that consists of several lines of defence. The first layer is the plant’s passive defence mechanism. These include structural barriers, such as the cell wall that can physically block the entry of pathogens into the plant tissues (Huckelhoven, 2007; Miedes et al., 2014). If pathogens are successful to pass a plant’s passive defences mechanism, they have the chance to access the nutrients from the plant. However, after the passive defence mechanism, plants still possess two layers of actively induced immune systems. The first layer of the immune response, which is activated by a pathogen-associated molecular pattern (PAMP), is PAMP-triggered immunity (PTI). PAMPs are a diverse set of microbial molecules that have conserved structures perceived by plant surface-exposed receptors called pattern recognition receptors (PRRs) (Medzhitov and Janeway, 1997; Macho and Zipfel, 2014). These membrane-bound PRRs are receptor-like kinases (RLK) or receptor-like proteins (RLP) with a high variety of intracellular domains. After the perception of PAMPs through PRRs, PAMP-triggered immunity (PTI) will be activated as the first layer of the plant immunity system or surface immunity, which restricts pathogen proliferation. PTI signaling components are often targeted by various pathogen virulence effector proteins, resulting in diminished plant defences and increased pathogen virulence. While PAMPs are conserved molecules that are shared among many different pathogens, effectors are typically species, race, or strain-specific molecules that contribute to pathogen virulence by targeting specific host plant processes (Thomma et al., 2011). Some plant resistance (R) proteins have evolved to recognize pathogen effectors directly or indirectly by associating with cytoplasmic immune receptors (Zhang J. and Zhou JM., 2010). These receptors, which often contain a nucleotide-binding leucine-rich repeat (NB-LRR) domain, activate the second layer of immunity known as effector-triggered immunity (ETI) (Macho and Zipfel, 2014). In contrast to PTI, ETI is a highly specific defense response that is triggered by the recognition of pathogen effectors that have been specifically adapted to interact with and manipulate host proteins. It can trigger programmed cell death, called the hypersensitive response (HR), in response to a pathogen attack, which helps to contain the infection and limit the damage caused to the plant (Coll et al., 2011).

The most effective management strategy to control SCN populations is using resistant cultivars rather than any other strategies, such as crop rotation or nematicides. Three popular Plant Introductions (PIs) resistant to SCN are PI 437654, PI 548402 (a.k.a. Peking) and PI 88788, which carry resistant loci effective against multiple nematode races (Brucker et al., 2005; Concibido et al., 2004). However, among all these resistant lines, plant breeders have heavily overused PI 88788 in the past three decades as resistant parent in breeding programs. This has led to the selection of virulent biotypes of SCN and population shifting such as HG type 1.2.5.7, which are able to overcome the PI 88788-type resistance.

Soybean cyst resistance is a complex trait with polygenic inheritance. The first quantitative trait loci (QTL) underlying the resistance to *H. glycines* (i.e., rhg) were reported in the early 1960s (Caldwell et al., 1960; Matson & Williams, 1965). Among several reported QTL, the QTL on chromosomes 18 (*rhg1*) and 8 (*Rhg4*) are the two major resistance ones that have been consistently mapped and reported in a variety of soybean germplasm (Concibido et al., 2004; Kim et al. 2010; Lu et al. 2022). In some SCN-resistant lines, such as PI 88788, *rhg1* with recessive action is sufficient to provide resistance to certain races of SCN and display an incompatible interaction with the nematode (Concibido et al., 2004). In other resistant sources, such as PI 548402, resistance to SCN requires both *rhg1* and *Rhg4,* while *Rhg4* exhibits a dominant gene action (Meksem et al., 2001). Brucker et al. 2005 classified *rhg1* into two types: *rhg1-a* in PI 548402 (a.k.a. Peking-type) with low copy number (three or fewer repeats), which reacts with *Rhg4* to provide greater resistance to SCN, and *rhg1-b* in PI 88788-type soybeans, which poses high copy number (four or more repeats) and provides the resistance without interacting with *Rhg4* (Brucker et al., 2005; Cook et al., 2012). SCN-susceptible lines such as Williams 82 and Lee74 have only a single copy of *rhg1* (Cook et al., 2012). Previous studies discovered the SCN resistance governed by *rhg1* is mediated through a 31-kb segment that is tandemly repeated and carries three genes, including a predicted amino acid transporter (*Glyma18g02580*), an SNAP protein predicted to participate in the disassembly of SNARE membrane trafficking complexes (*Glyma18g02590*), and a protein with aWI12 (wound-inducible protein 12) region without functionally characterized domains (*Glyma18g02610*) (Cook et al., 2012, 2014). On the other hand, map-based cloning of the *Rhg4* locus revealed that a single gene encoding a serine hydroxymethyltransferase (SHMT, *Glyma08g11490*) is responsible for the resistance (Liu et al., 2012).

By deploying defense mechanisms, SCN-resistant soybean genotypes are able to mount effective immune responses against SCN and minimize the damage caused to their production. However, pathogens have also evolved sophisticated strategies to evade or overcome these defenses, which has led to an ongoing evolutionary arms race between plants and pathogens. Therefore, studying soybean-SCN interactions can be challenging due to the complex nature of these interactions and the fact that the molecular mechanisms involved can be highly dynamic and context-dependent. It is important to use sensitive and high-throughput methods that can capture the complexity of these interactions and provide a comprehensive view of the molecular changes that occur during infection.

RNA sequencing enables high throughput analysis of the transcriptome landscape of cells. In host cells infected with pathogens, two organisms interact with markedly distinct transcriptomes. Dual RNA sequencing is a powerful in-silico analysis method that enables the simultaneous study of the gene expression responses of both pathogens and host cells from the same samples, thereby deepening our understanding of their interaction (Westermann et al., 2016; Mika-Gospodorz et al., 2020). In this study, we aimed to understand how SCN invasion modulates soybean gene expression, while simultaneously examining pathogen reactions across multiple hosts (e.g. compatible, semi-compatible, semi-incompatible, incompatible soybean). By utilizing dual RNA sequencing, we provide novel insights into the intricate interplay between host and parasite, revealing a diversity of defense mechanisms in soybean and virulence genes in the soybean cyst nematode. This study marks the first time that such a method has been employed to investigate soybean-SCN interactions, offering new insights into this crucial area of research.

## Results

### Greenhouse SCN bioassay and genotyping *rhg1* and *Rhg4* copy number variation (CNV)

The tolerance of the lines PI 88788, PI 437654, PI 548402 and Lee 74 were characterized to their level of resistance according to the scaled proposed by Schmitt and Shannon 1992 (Table 1). The average number of females on Lee74 was 110.2 and the highest to the lowest FI value was 63.0, 10.0, and 0.0 on PI 88788, PI 548402, and PI 437654, respectively (Figure 1). However, despite high levels of SCN replication on PI 88788 following inoculation with SCN HG type 1.2.5.7, the egg production was significantly lower than that of Lee74 (Figure1). Considering these results, in this study we called PI 437654 as a incompatible line, PI 548402 as a semi-incompatible line, PI 88788 as a semi-compatible line and Lee 74 as a compatible line.

**Table 1.** Female index scale of tested soybean lines against SCN HG type 1.2.5.7

| Female index (%) | Rating | Label |
| --- | --- | --- |
| <10 | Resistant | R |
| ≥10 to 30 | Moderately resistant | MR |
| > 30 to 60 | Moderately susceptible | MS |
| > 60 | Susceptible | S |

**Figure 1.**
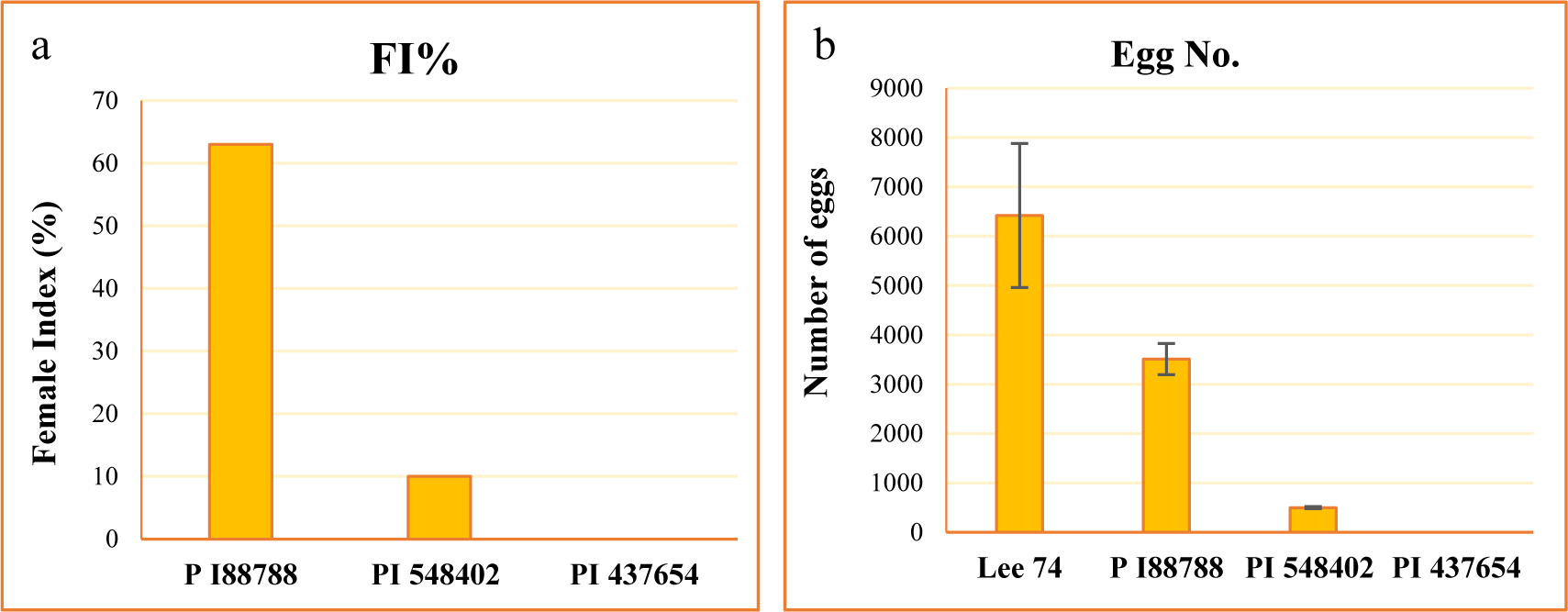
Estimation of FI values (a) and counting number of eggs (b) in each line. Standard error (SE) at α = 0.05.

Based on the results mentioned above, which include the FI values and the number of eggs in the tested PI lines, we decided to focus primarily on the PI 437654 defense mechanism. This line has shown complete resistance against SCN HG-type 1.2.5.7. To validate the SCN responsive genes of PI 437654, we utilized PI 548402 and PI 88788, which have strong and weak defense mechanisms, respectively, against SCN HG type 1.2.5.7.

We evaluated the copy number variation (CNV) of the *rhg1* and *Rhg4* genes, well-studied loci conferring resistance to SCN, in all of the tested PI lines. To do this, we developed a Taqman assay from the conserved regions of these genes. The assay results indicated that the CNV of the *rhg1* locus was 8, 3, 3, and 1 in PI 88788, PI 548402, PI 437654 and Lee74, respectively (Figure 2a). Additionally, the CNV of *Rhg4* was 1, 2, 3, and 1 in these same lines, respectively (Figure 2b).

**Figure 2.**
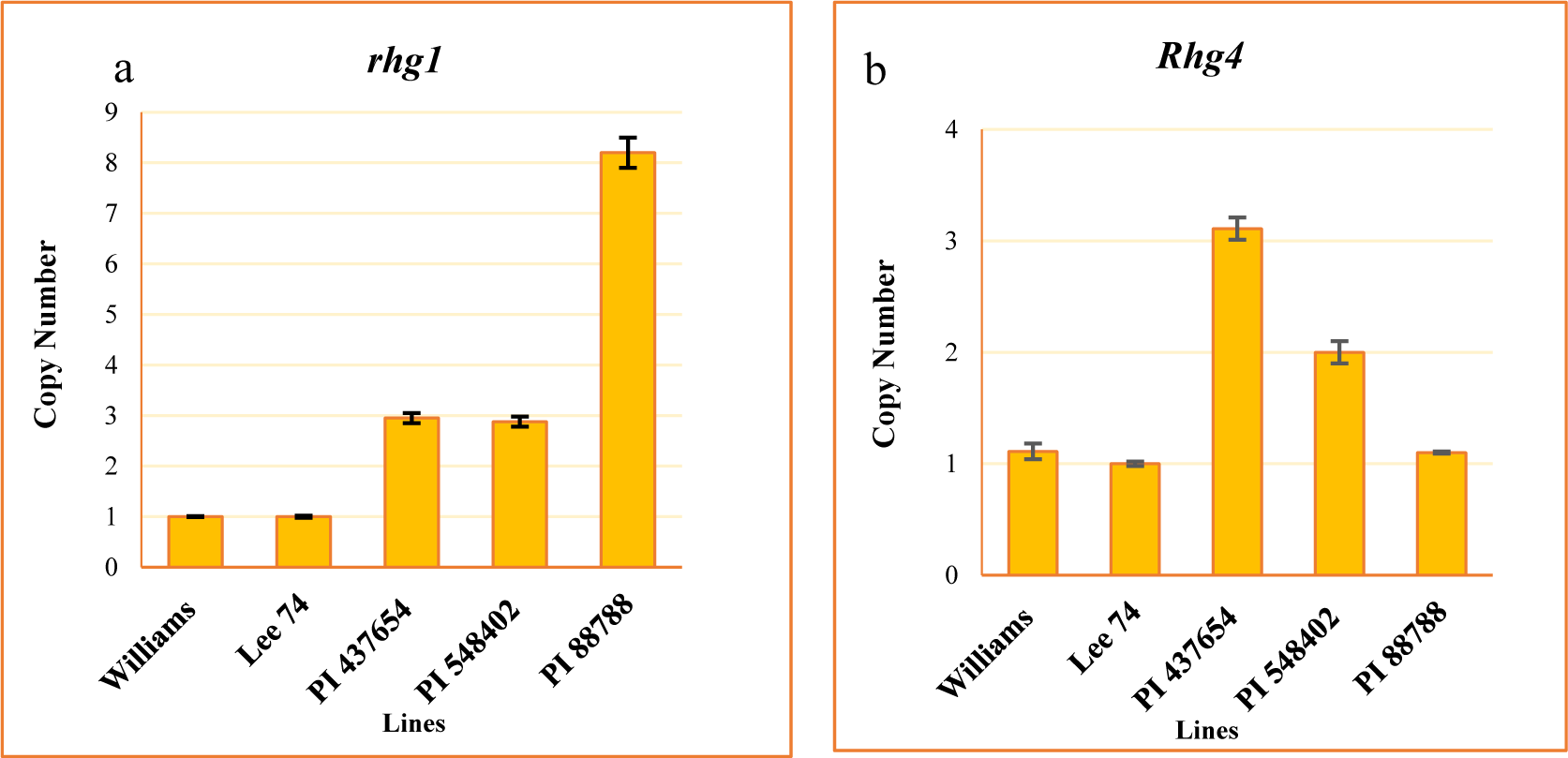
CNV of *rhg1* (a) and *Rhg4* (b). Williams used as a calibrator. Standard error (SE) at α = 0.05.

### Dual transcriptome sequencing and assembly in host and nematode

We conducted a comprehensive analysis of the transcriptomic response of soybean to SCN, along with the reaction of SCN to different soybean hosts. RNA sequencing was performed on the four soybean PI lines, the three resistant lines (PI 88788, PI 548402, and PI 437654) and the susceptible line (Lee74) at 5 and 10 days post-inoculation (dpi) with SCN HG type 1.2.5.7. Prior to conducting RNA sequencing, we confirmed infection of the roots with SCN HG type 1.2.5.7 at 5 and 10 dpi by amplifying the SCN 18S ribosomal gene from DNA extracted from the roots (Figure S1). Global patterns of gene expression in both soybean and *H. glycine* were evaluated using high-throughput RNA sequencing, with each sample having a high coverage of at least 50 million reads. In total, we obtained 58 million reads per sample from both infested and control conditions, of which approximately 87.83% were successfully mapped to the soybean reference transcriptome (Table S1). When examining SCN HG type 1.2.5.7 at 5 dpi, we found that Lee74 had the highest total number of reads mapped to *H. glycine*, while PI 437654 had the lowest (Figure 3, Table S1). At 10 dpi, the total number of reads mapped to *H. glycine* had significantly decreased in Lee74, PI 548402, and PI 88788. The highest number of reads was obtained from PI 88788 (Figure 3).

**Figure 3.**
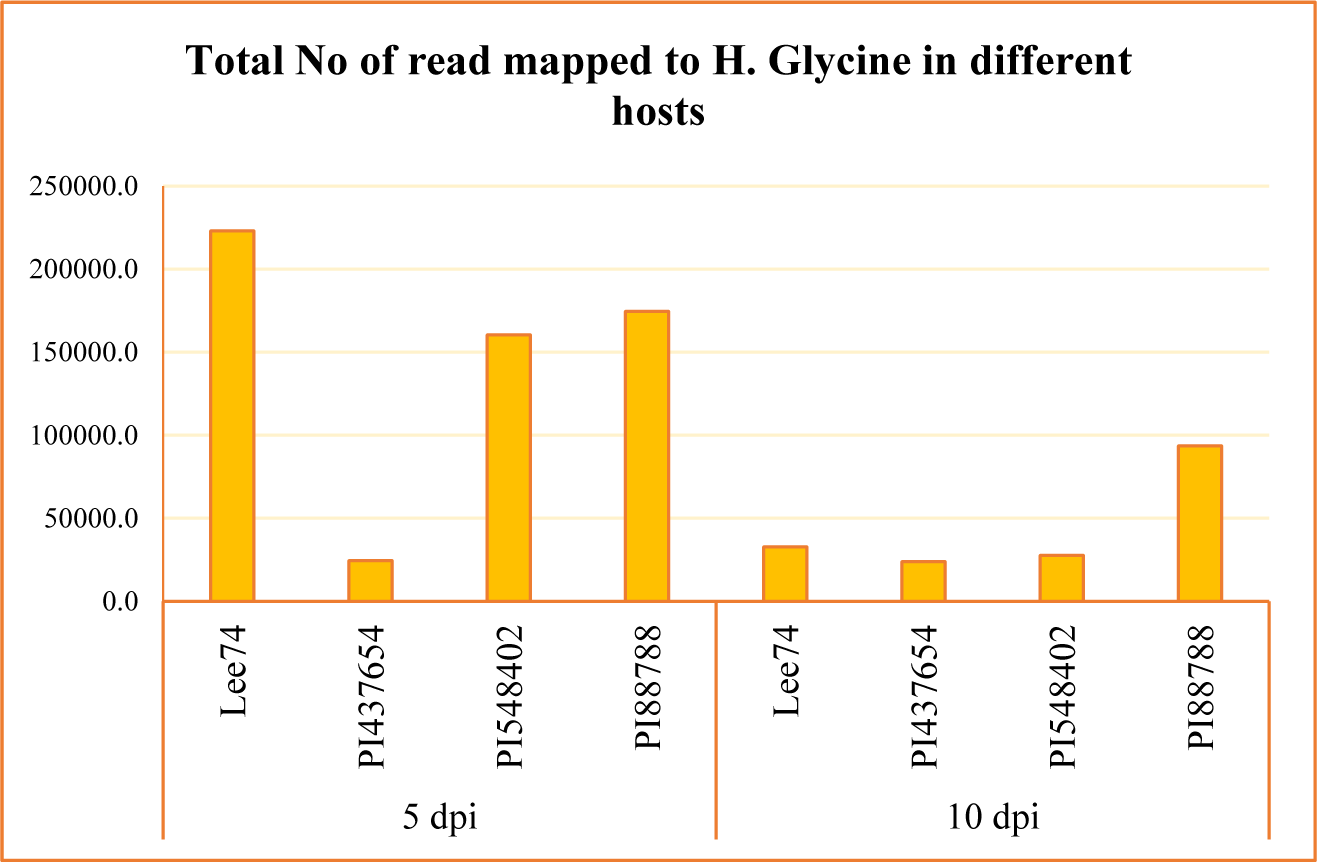
Total number of read mapped to *H. Glycine* in four different hosts (e.g., Lee74, PI 437654, PI 548402 and PI 88788) and in two time points (e.g., 5 & 10 dpi)

### Intra- and Inter-genotype analyses identify pathways pertaining to defence in each resistant line

When examining differentially expressed genes (DEGs) within each genotype under infested and non-infested conditions, we identified that 167, 1966, 581, 161 genes (DEGs) were upregulated (with cut-offs of log_2_ fold change ≥ 1, fragments per kilobase (kb) of transcript per million mapped reads (FPKM) ≥ 1, and p < 0.05) in PI 437654, PI 548402, PI 88788 and Lee74, respectively, at 5 dpi (Figure 4). At 10 dpi, the number of DEGs significantly decreased in PI 437654, PI 548402, and PI 88788 but the number of DEGs remarkably increase in Lee74, (Figure 4).

**Figure 4.**
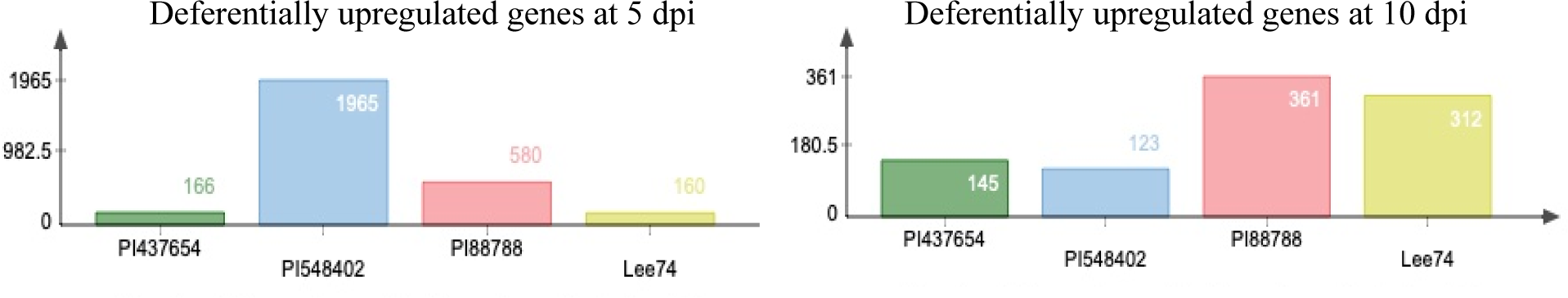
The reaction or differentially upregulated genes in PI 88788, PI 548402, PI 437654 and Lee74 against SCN HG type 1.2.5.7

Intra-genotype gene ontology enrichment (GOE) analyses revealed functional categories, defined by high-level GO terms, that are involved in the immunity response of each line. In PI 437654, 44 out of 166 upregulated genes were found to be involved in the over-represented GO terms at 5 dpi (Table 2). Based on GO biological process analyses, these upregulated genes in PI 437654 at 5 dpi clustered into four groups: (1) programmed cell death in response to reactive oxygen species (ROS), (2) cell detoxification or response to stress, (3) carbon fixation, and (4) salicylic acid biosynthesis (Table S2). At 10 dpi, 65 genes out of 145 upregulated genes were assigned to over-represented GO terms (Table 3). These genes clustered in three groups: (1) detoxification or response to stress, (2) salicylic acid biosynthesis, and (4) sulfur and glycosyl metabolic process (Table S3).

**Table 2.**
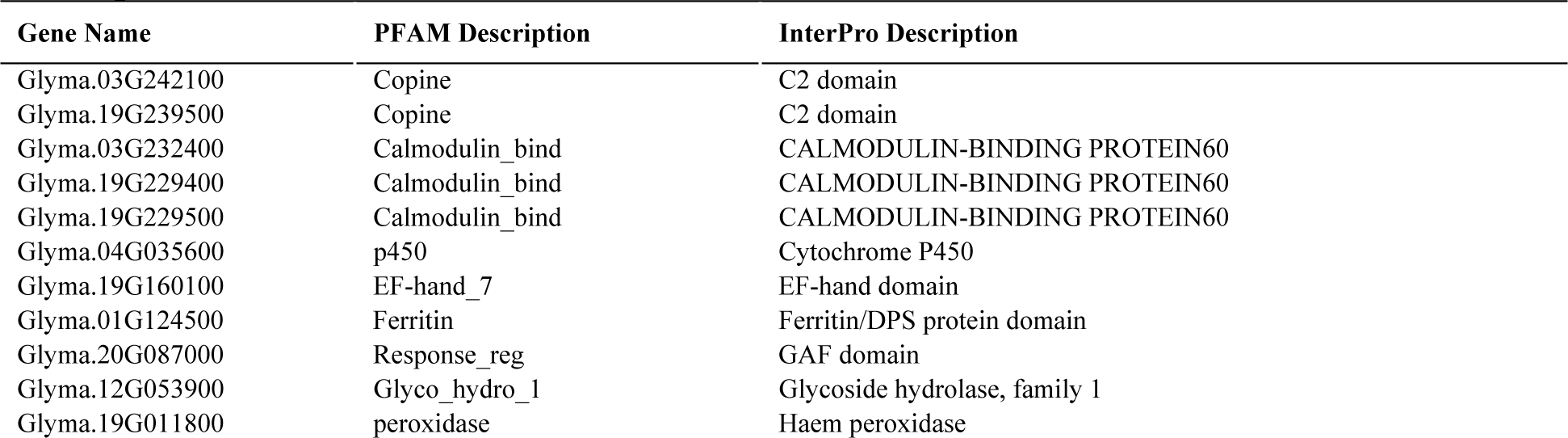
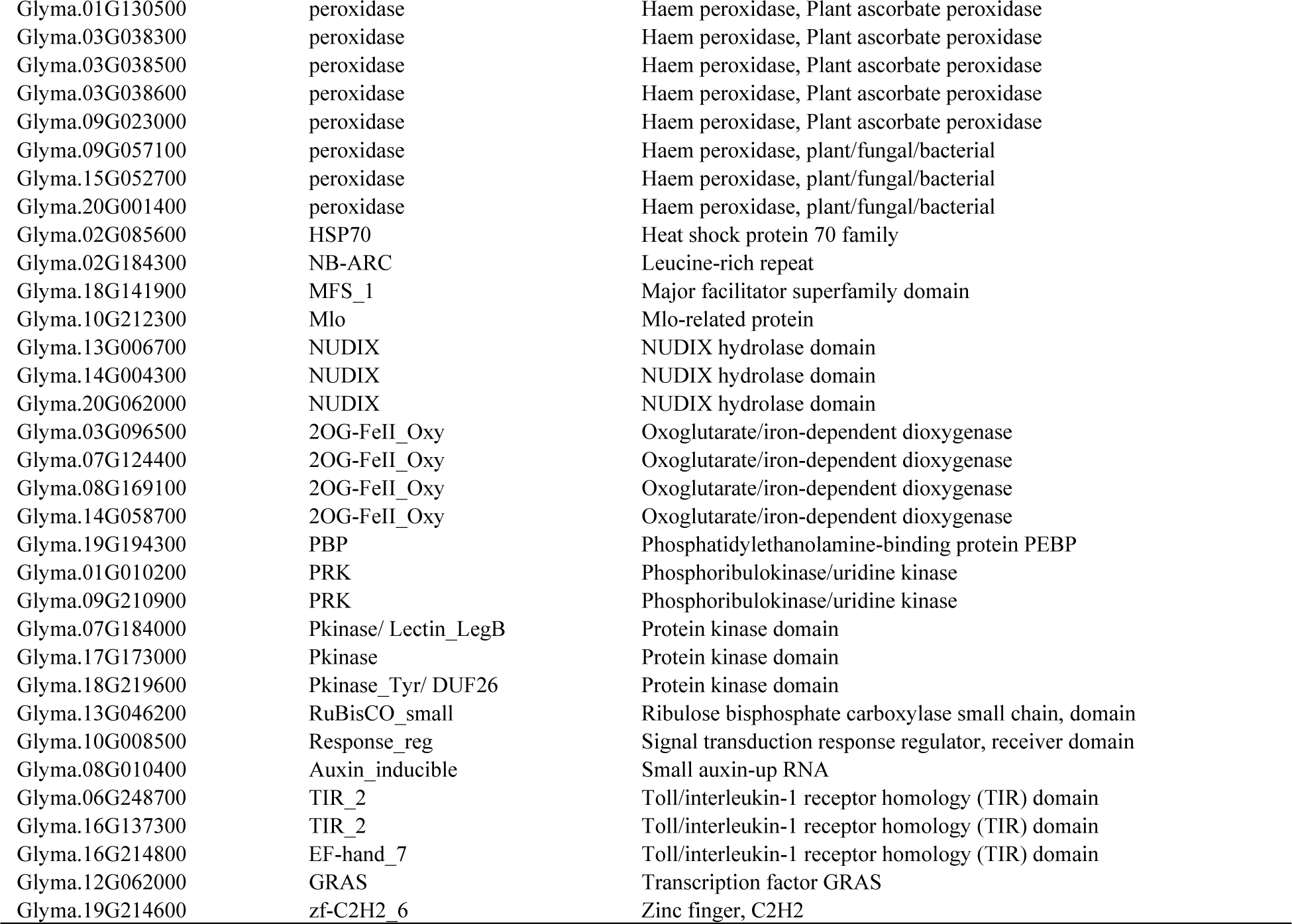
PI 437654 upregulated genes that were assigned in over-represented GO terms <u>in 5 dpi</u>

| Gene Name | PFAM Description | InterPro Description |
| --- | --- | --- |
| Glyma.03G242100 | Copine | C2 domain |
| Glyma.19G239500 | Copine | C2 domain |
| Glyma.03G232400 | Calmodulin_bind | CALMODULIN-BINDING PROTEIN60 |
| Glyma.19G229400 | Calmodulin_bind | CALMODULIN-BINDING PROTEIN60 |
| Glyma.19G229500 | Calmodulin_bind | CALMODULIN-BINDING PROTEIN60 |
| Glyma.04G035600 | p450 | Cytochrome P450 |
| Glyma.19G160100 | EF-hand_7 | EF-hand domain |
| Glyma.01G124500 | Ferritin | Ferritin/DPS protein domain |
| Glyma.20G087000 | Response_reg | GAF domain |
| Glyma.12G053900 | Glyco_hydro_1 | Glycoside hydrolase, family 1 |
| Glyma.19G011800 | peroxidase | Haem peroxidase |
| Glyma.01G130500 | peroxidase | Haem peroxidase, Plant ascorbate peroxidase |
| Glyma.03G038300 | peroxidase | Haem peroxidase, Plant ascorbate peroxidase |
| Glyma.03G038500 | peroxidase | Haem peroxidase, Plant ascorbate peroxidase |
| Glyma.03G038600 | peroxidase | Haem peroxidase, Plant ascorbate peroxidase |
| Glyma.09G023000 | peroxidase | Haem peroxidase, Plant ascorbate peroxidase |
| Glyma.09G057100 | peroxidase | Haem peroxidase, plant/fungal/bacterial |
| Glyma.15G052700 | peroxidase | Haem peroxidase, plant/fungal/bacterial |
| Glyma.20G001400 | peroxidase | Haem peroxidase, plant/fungal/bacterial |
| Glyma.02G085600 | HSP70 | Heat shock protein 70 family |
| Glyma.02G184300 | NB-ARC | Leucine-rich repeat |
| Glyma.18G141900 | MFS_1 | Major facilitator superfamily domain |
| Glyma.10G212300 | Mlo | Mlo-related protein |
| Glyma.13G006700 | NUDIX | NUDIX hydrolase domain |
| Glyma.14G004300 | NUDIX | NUDIX hydrolase domain |
| Glyma.20G062000 | NUDIX | NUDIX hydrolase domain |
| Glyma.03G096500 | 2OG-FeII_Oxy | Oxoglutarate/iron-dependent dioxygenase |
| Glyma.07G124400 | 2OG-FeII_Oxy | Oxoglutarate/iron-dependent dioxygenase |
| Glyma.08G169100 | 2OG-FeII_Oxy | Oxoglutarate/iron-dependent dioxygenase |
| Glyma.14G058700 | 2OG-FeII_Oxy | Oxoglutarate/iron-dependent dioxygenase |
| Glyma.19G194300 | PBP | Phosphatidylethanolamine-binding protein PEBP |
| Glyma.01G010200 | PRK | Phosphoribulokinase/uridine kinase |
| Glyma.09G210900 | PRK | Phosphoribulokinase/uridine kinase |
| Glyma.07G184000 | Pkinase/ Lectin LegB | Protein kinase domain |
| Glyma.17G173000 | Pkinase | Protein kinase domain |
| Glyma.18G219600 | Pkinase_Tyr/ DUF26 | Protein kinase domain |
| Glyma.13G046200 | RuBisCO_small | Ribulose biphosphate carboxylase small chain, domain |
| Glyma.10G008500 | Response_reg | Signal transduction response regulator, receiver domain |
| Glyma.08G010400 | Auxin_inducible | Small auxin-up RNA |
| Glyma.06G248700 | TIR_2 | Toll/interleukin-1 receptor homology (TIR) domain |
| Glyma.16G137300 | TIR_2 | Toll/interleukin-1 receptor homology (TIR) domain |
| Glyma.16G214800 | EF-hand_7 | Toll/interleukin-1 receptor homology (TIR) domain |
| Glyma.12G062000 | GRAS | Transcription factor GRAS |
| Glyma.19G214600 | zf-C2H2_6 | Zinc finger, C2H2 |

**Table 3.**
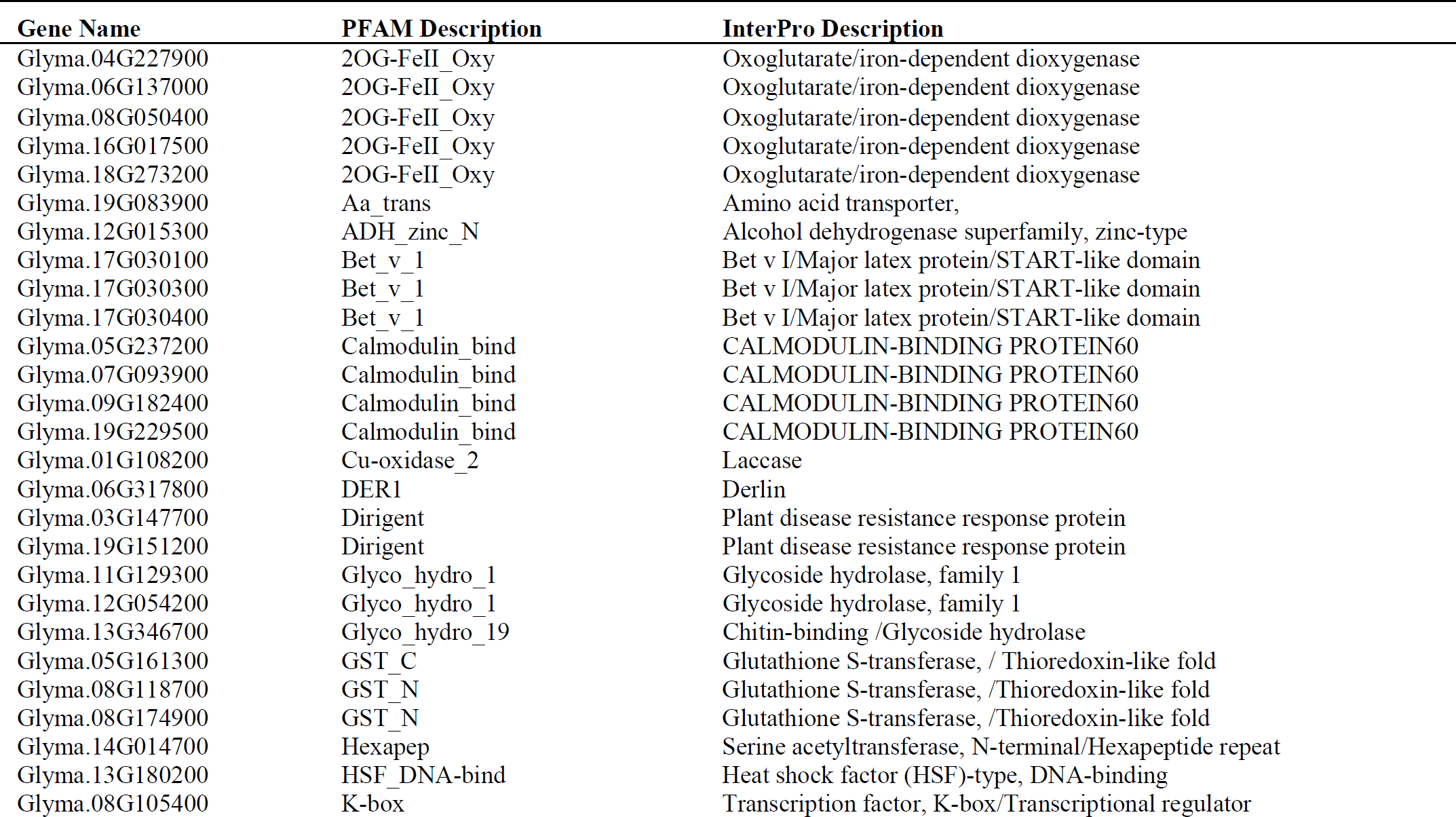
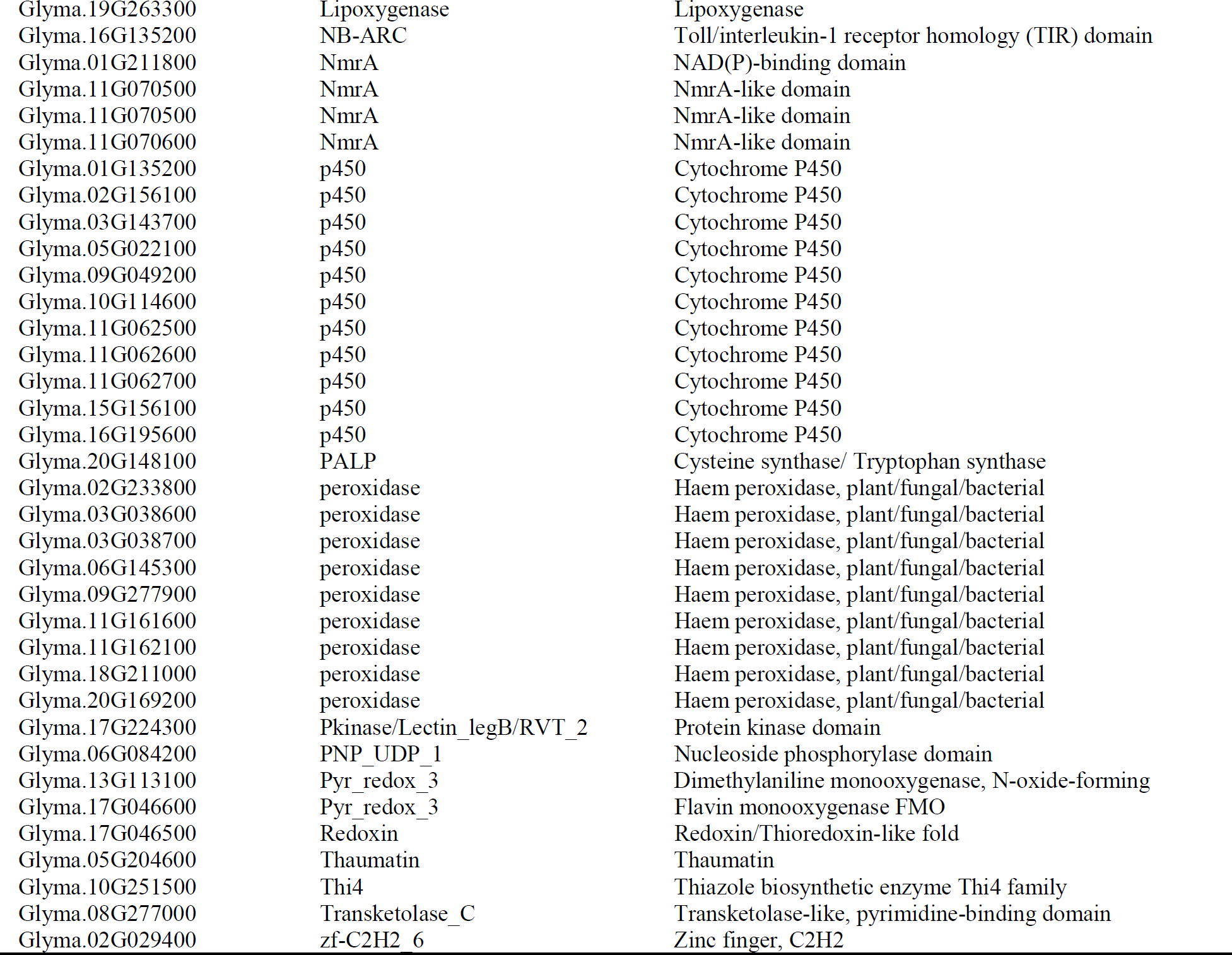
PI 437654 upregulated genes that were assigned in over-represented GO terms <u>in 10 dpi</u>

Inter-genotype analyses of DEGs that were significantly upregulated in all lines at 5 dpi revealed that PI 437654, a strong source of resistance (FI=0%), has 63 unique genes that were not differentially expressed in other resistant lines, PI 548402, PI 88788, with lower levels of resistance, FI=10.0% and 63.0, respectively, or in Lee74, the susceptible line (Figure 5). In addition, 42 SCN-responsive genes were detected in PI 437654, which were also found to be upregulated in PI 548402 (Figure 5). Comparison of PI 437654 with PI 548402 and PI 88788 resulted in the identification of 44 differentially upregulated genes that overlapped among these three resistant lines (Figure 5). Although our focus was on PI 437654, through this comparison we validated those SCN-responsive genes that were also observed in PI 548402 or/and PI 88788.

**Figure 5.**
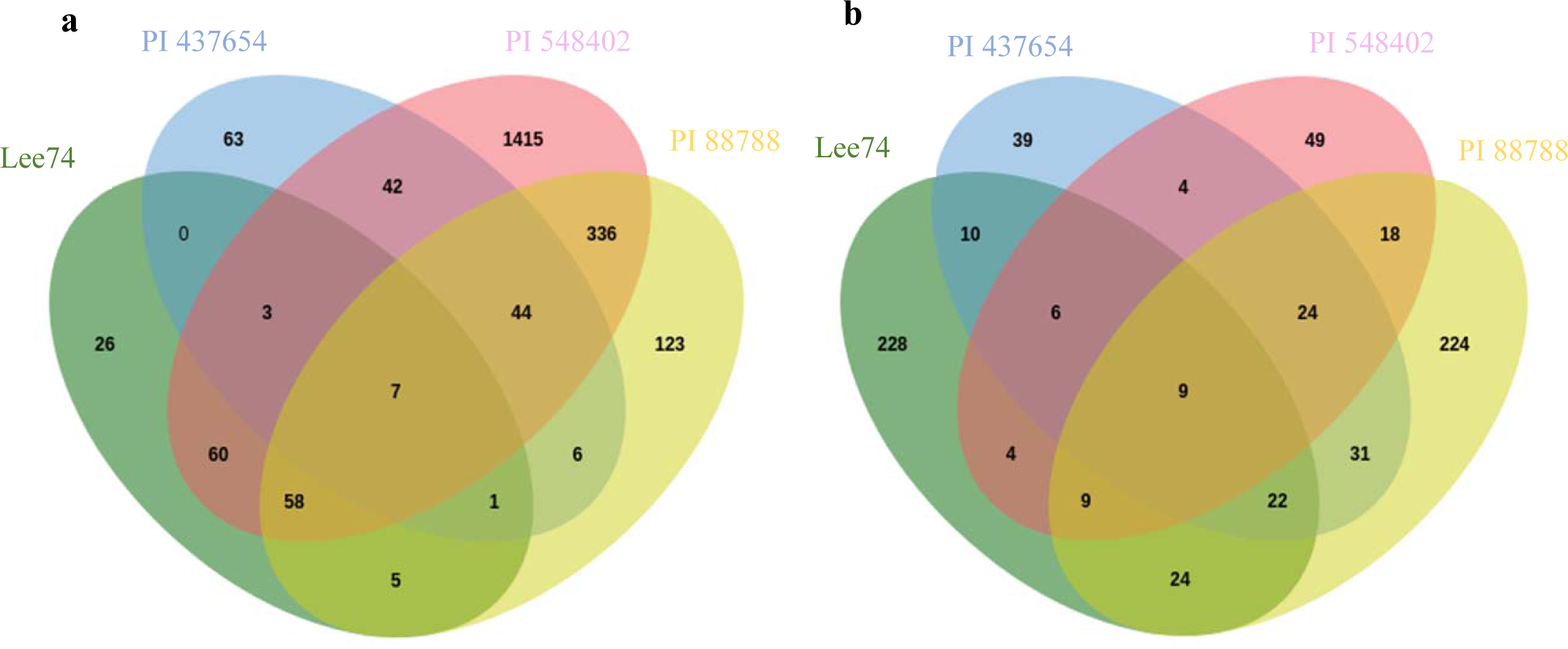
Venn diagram showing the number of unique and overlapping SCN responsive sets of genes in PI 437654 compared to PI 548402, PI 88788, and Lee74 which were upregulated at 5 dpi (a) and 10 dpi (b)

To gain insight into the functional categories of genes that are differentially expressed in response to soybean cyst nematode (SCN) infection, we conducted GO enrichment analyses on uniquely differentially expressed genes in PI 437654, as well as on those genes that overlapped with PI 548402 and PI 88788 (Tables 4 & 5). We identified several over-represented GO term categories and corresponding genes involved in these categories at 5 dpi. In PI 437654, six GO term categories were identified among the 63 uniquely upregulated genes (Figure 5), including response to chemical, negative regulation of development and reproductive process, photosynthesis and carbon fixation, salicylic acid metabolic process, oxazole or thiazole metabolic process, and fructose1,6 bisphosphate metabolic process (Table S4). At the same time point, 16 genes out of 42 significantly upregulated genes detected in both PI 437654 and PI 548402 were found to be enriched in four distinct GO terms, including response to chemical and oxidative stress, programmed cell death in response to ROS, cellular response to an inorganic substance, and response to ethylene (Table 4, Table S5 and Figure 5). In addition, we identified 20 SCN-responsive genes overlapped in PI 437654, PI 548402 and PI 88788 and categorized them into two over-represented GO terms: response to stress & cellular detoxification, and regulation of primary metabolic processes such as RNA and DNA (Table 4, Table S6 & Figure 5). Finally, we found that among the six common DEGs detected in both PI 437654 and PI 88788 at 5 dpi, only one gene (*Glyma_02G028000*) was categorized into an over-represented GO, specifically the Collagen catabolic process (Table 4 and Figure 5). These results provide valuable insights into the functional categories of genes that are involved in the soybean response to SCN infection and can help inform future studies aimed at improving soybean resistance to SCN.

**Table 4.**
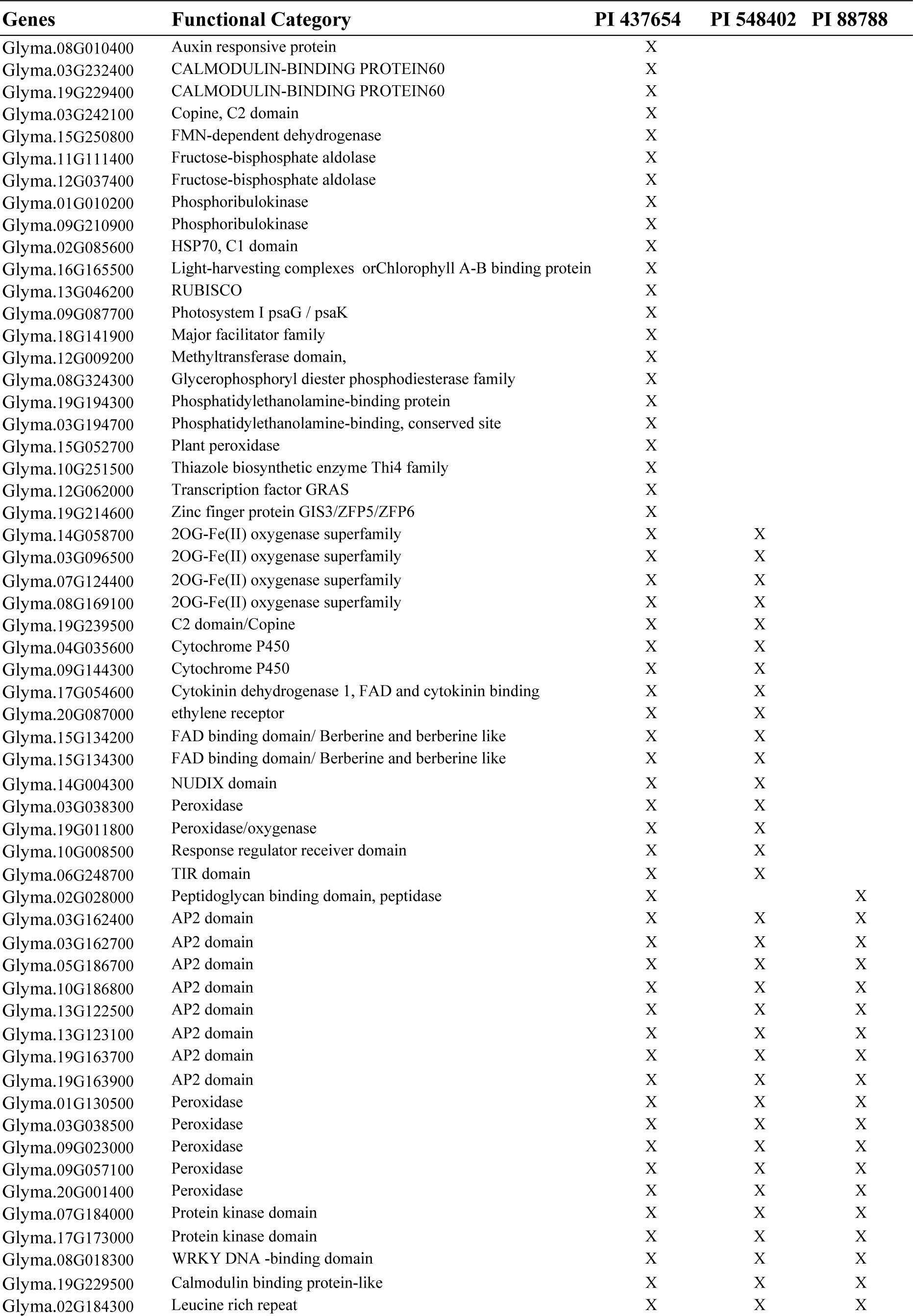
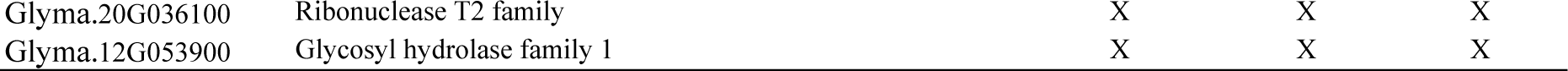
Sets of genes in over-represented GO terms of PI 437654 that were unique DEGs or validated by PI <u>548402 and/or PI 88788 at 5 dpi</u>

At 10 dpi, inter-genotype analyses revealed PI 437654 as a highly resistant source (FI=0%), displaying 39 unique genes not differentially expressed in other resistant lines, including PI 548402 and PI 88788, which showed lower resistance levels with FIs of 10.0% and 63.0%, respectively, and the susceptible line, Lee74 (Figure 5). Additionally, four SCN-responsive genes were identified in PI 437654 that were also upregulated in PI 548402, while 31 SCN-responsive genes were upregulated in PI 88788 (Figure 5). By comparing PI 437654 with PI 548402 and PI 88788, we identified 24 differentially upregulated genes that were shared among all three lines (Figure 5). This comparison allowed for validation of a portion of the SCN-responsive genes identified in PI 437654, using PI 548402 and PI 88788 as additional reference points.

At 10 dpi, GO enrichment analysis identified four enriched genes out of 39 unique DEGs in PI 437654, which were annotated in the GO terms ammonium transmembrane transport & sulfur amino acid biosynthetic process (Table 5 & Table S7). Furthermore, among the four common genes shared between PI 437654 and PI 548402, two were identified as part of a set of genes enriched in GO terms related to oxazole or thiazole biosynthetic process, S-glycoside catabolic process, and sulfur compound metabolic process (Table S8). GO enrichment analyses identified 14 genes out of 31 SCN-responsive genes that overlapped in PI 437654 and PI 88788 (Table 5 & Figure 5), which were categorized into five over-represented GO terms: salicylic acid biosynthetic process, monocarboxylic acid biosynthetic process, cellular oxidant detoxification, organic acid metabolic process, regulation of nucleic acid-templated transcription, and response to stress (Table S9). Among PI 437654, PI 548402, and PI 88788, there were 24 overlapped genes that we were not able to assign to any enriched GO terms (Table 5).

**Table 5.**
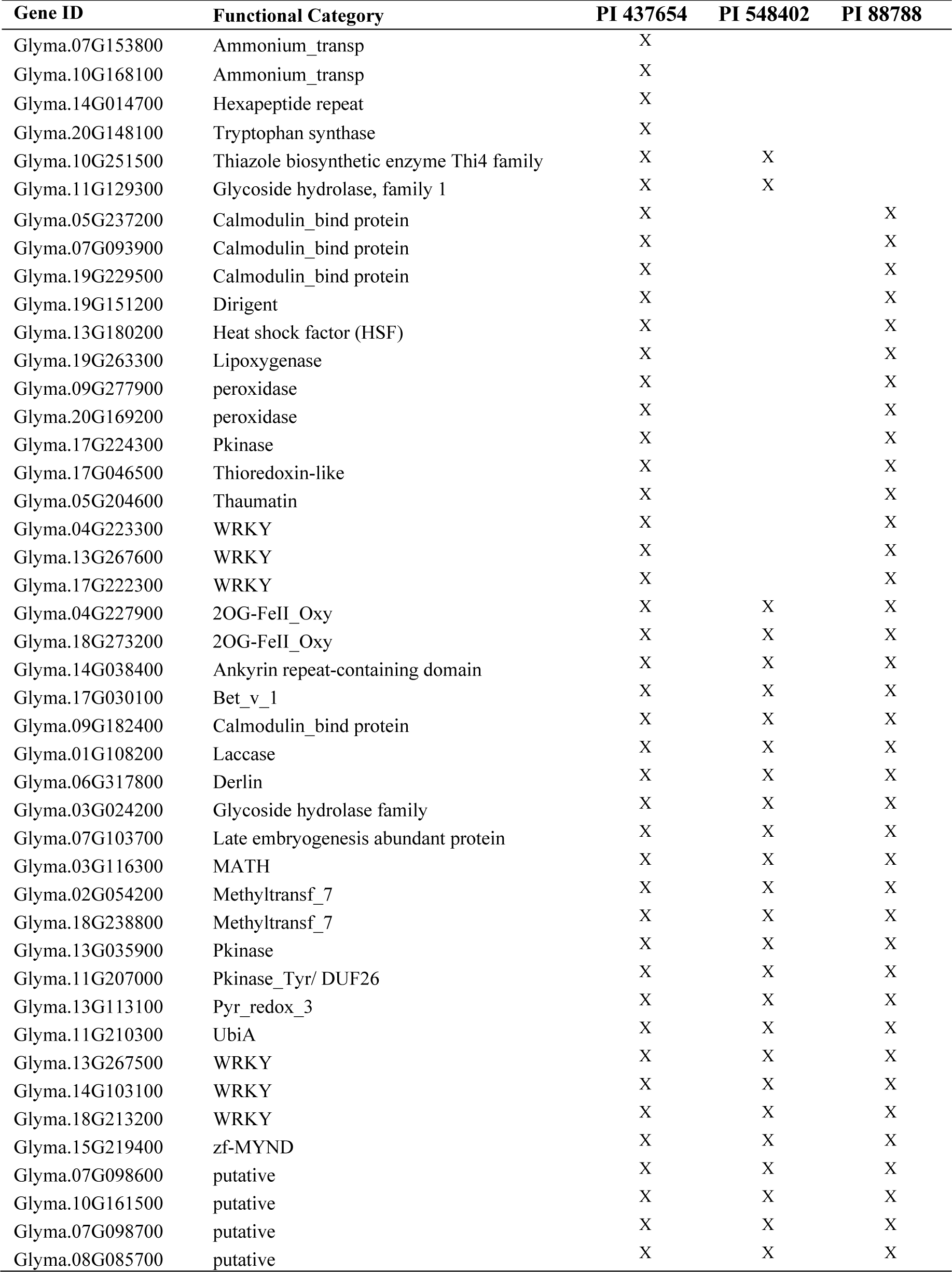
Sets of the genes in over-represented GO terms of PI 437654 that were unique DEGs or validated by PI548402 and/or PI88788 at 10 dpi

The pathway analysis of DEGs in PI 437654, PI 548402, PI 88788 and Lee74 revealed enriched immune response pathways in resistant lines. KEGG analysis indicated that the MAPK signaling pathway, phenylpropanoid biosynthesis, plant hormone signal transduction, and biosynthesis of secondary metabolite were significant pathways associated with soybean immune response to SCN HG Type 1.2.5.7 in PI 437654, PI 548402, and PI 88788 at 5 dpi (Table S10). In PI 437654, we observed that there are other pathways associated with the immune systems, such as plant-pathogen interaction, carbon fixation in the photosynthetic organism, and carbon metabolism. These pathways set PI 437654 apart from PI 548402 and PI 88788, making it a unique SCN source (Table S10). Pathway analysis of PI 437654 at 10 dpi revealed that upregulated DEGs were enriched in several metabolic pathways, including phenylpropanoid biosynthesis, isoflavonoid biosynthesis, flavonoid biosynthesis, thiamine metabolism, glutathione metabolism, sulfur metabolism, cysteine and methionine metabolism, cyanoamino acid metabolism, circadian rhythm, biosynthesis of cofactors, biosynthesis of secondary metabolites, ubiquinone and terpenoid-quinone biosynthesis (Table S11).

### Interactions between SCN transcriptomes and soybean proteins induces either susceptibility or resistance

In order to study the expression of SCN pathogenicity genes during the infection process and assess the behaviour of SCN towards different hosts, we analyzed the transcriptomes of SCN HG type 1.2.5.7 at two-time points in the four soybean PI lines. Comparison of the SCN transcriptomes across different soybean lines showed that the pathogenicity gene expression can be altered by the host plant. Furthermore, a two-by-two comparison of the host plants revealed that the SCN HG type 1.2.5.7 transcriptome in interaction with Lee 74 exhibited the largest number of differentially expressed genes compared with the transcriptome in PI 437654. In contrast, the transcriptome of SCN HG type 1.2.5.7 in interaction with Lee 74 had the highest similarity in gene expression profile with PI 88788 (Figure 6). For instance, the SCN transcriptome in Lee 74 had 123 differentially upregulated and 136 differentially downregulated genes when compared with its transcriptome in PI 437654, while only one gene was differentially upregulated in Lee 74 when compared with PI 88788 (Figure 6). This suggests that SCN adapts its gene expression profile in response to the host plant and that different soybean lines may have different levels of resistance against SCN.

**Figure 6.**
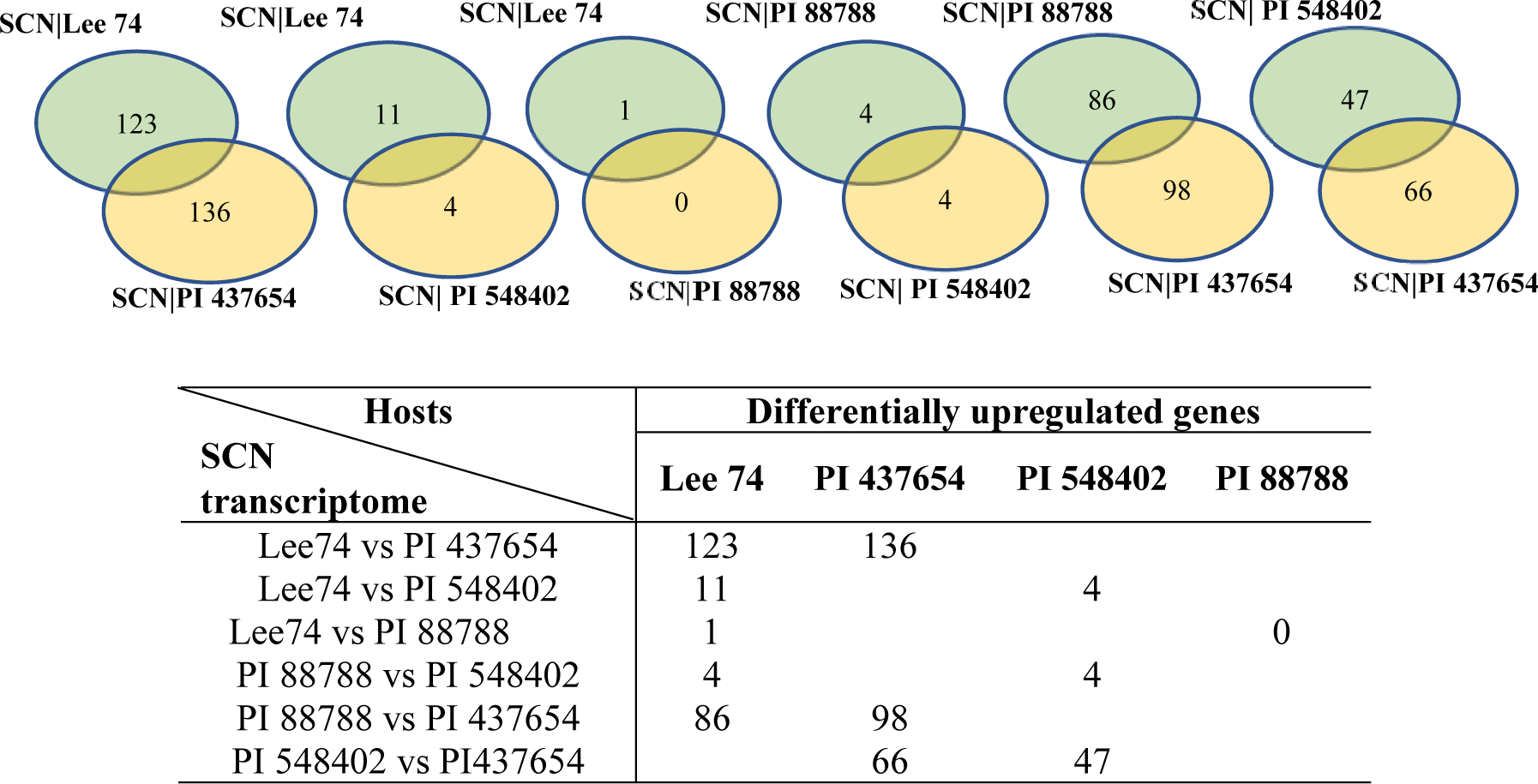
Number of genes differentially express in SCN HG type 1.2.5.7 transcriptome in reaction with the four different hosts

Effector genes play a crucial role in the molecular interactions between SCN and soybean. These genes encode secretory proteins that are translocated into the root tissue of soybean during infection. In this study, putative effectors were identified in four soybean lines (PI 437654, PI 548402, PI 88788, and Lee 74). Analysis of putative effectors revealed that PI 437654 had a unique set of effectors, unlike the other soybean hosts. Furthermore, the putative effectors of PI 548402 shared a 62% similarity with those of PI 88788 and 50% similarity with Lee 74. The most commonly shared effector genes were found between PI 88788 and Lee 74, with 89% similarity (Table 6). These findings highlight the importance of effectors in the pathogenesis of *H. glycines* and provide insight into the diversity and conservation of these genes in different soybean hosts.

**Table 6.** Differentially expressed SCN putative effector genes in different soybean lines: PI437654, PI 548402, PI88788, Lee74

| Identified Effector |  | Probability of Predicted signals & localizations |  | Chr | Significantly Upregulated |  |  |  | Reported Effectors |
| --- | --- | --- | --- | --- | --- | --- | --- | --- | --- |
| Genome draft format-ID | Pseudomolecule assembly-ID | Signal Peptide | Extra-cellular |  | PI437654 | PI 548402 | PI88788 | Lee74 |  |
| Hetgly.G000000386 | Hetgly10240 | 0.9556 | 0.8421 | 2 | X |  |  |  | NO |
| Hetgly.G000002041 | Hetgly00243 | 0.8547 | 0.5982 | 5 | X |  |  |  | NO |
| Hetgly.G000003725 | Hetgly01606 | 0.9997 | 0.9750 | 5 |  |  | X | X | NO |
| Hetgly.G000003742 | Hetgly01620 | 0.5363 | 0.7217 | 5 |  | X | X | X | NO |
| Hetgly.G000005495 | Hetgly01486 | 0.9995 | 0.9838 | 5 | X |  |  |  | NO |
| Hetgly.G000005767 | Hetgly10721 | 0.9997 | 0.5519 | 2 | X |  |  |  | NO |
| Hetgly.G000005859 | Hetgly07384 | 0.9996 | 0.9257 | 1 |  | X |  |  | NO |
| Hetgly.G000006034 | Hetgly15160 | 0.9996 | 0.7696 | 6 | X |  |  |  | NO |
| Hetgly.G000006271 | Hetgly13266 | 0.9951 | 0.9072 | 4 | X |  |  |  | YES<sup>1,2</sup> |
| Hetgly.G000007737 | Hetgly14373 | 0.9997 | 0.9553 | 6 | X |  |  |  | YES<sup>1</sup> |
| Hetgly.G000008328 | Hetgly08786 | 0.9998 | 0.9278 | 1 | X |  |  |  | YES<sup>1</sup> |
| Hetgly.G000008629 | Hetgly10435 | 0.8469 | 0.7313 | 2 |  |  | X |  | NO |
| Hetgly.G000008756 | Hetgly04908 | 0.9998 | 0.9513 | 3 |  | X | X | X | NO |
| Hetgly.G000008760 | Hetgly04892 | 0.9997 | 0.9781 | 3 | X |  |  |  | YES<sup>1</sup> |
| Hetgly.G000009320 | Hetgly14722 | 0.8276 | 0.9354 | 6 |  | X |  |  | NO |
| Hetgly.G000009584 | Hetgly20383 | 0.9997 | 0.9577 | 9 |  | X | X | X | NO |
| Hetgly.G000009600 | Hetgly20345 | 0.9997 | 0.9135 | 9 |  |  | X | X | NO |
| Hetgly.G000009601 | Hetgly20346 | 0.9992 | 0.8075 | 9 |  |  | X | X | NO |
| Hetgly.G000010445 | Hetgly05887 | 0.9648 | 0.9543 | 3 |  | X | X | X | NO |
| Hetgly.G000010619 | Hetgly08167 | 0.9997 | 0.9611 | 1 | X |  |  |  | YES<sup>1</sup> |
| Hetgly.G000011018 | Hetgly00611 | 0.9971 | 0.9794 | 5 | X |  |  |  | NO |
| Hetgly.G000011037 | Hetgly21117 | 0.9997 | 0.9685 | 8 |  | X | X | X | NO |
| Hetgly.G000011113 | Hetgly05749 | 0.9998 | 0.9162 | 3 |  |  |  | X | NO |
| Hetgly.G000011117 | Hetgly05752 | 0.9998 | 0.9662 | 3 |  | X |  |  | NO |
| Hetgly.G000011166 | Hetgly05791 | 0.9997 | 0.6696 | 3 | X |  |  |  | NO |
| Hetgly.G000011538 | Hetgly06927 | 0.7568 | 0.5704 | 1 | X |  |  |  | NO |
| Hetgly.G000011604 | Hetgly19536 | 0.6212 | 0.9683 | 9 |  | X | X |  | NO |
| Hetgly.G000011607 | Hetgly19535 | 0.9996 | 0.9769 | 9 |  | X | X | X | NO |
| Hetgly.G000012678 | Hetgly20828 | 0.9996 | 0.5010 | 8 | X |  |  |  | NO |
| Hetgly.G000013560 | Hetgly06640 | 0.9998 | 0.7309 | 1 |  |  | X | X | YES<sup>1</sup> |
| Hetgly.G000014327 | Hetgly13555 | 0.9998 | 0.9694 | 4 | X |  |  |  | YES<sup>1</sup> |
| Hetgly.G000014444 | Hetgly02677 | 0.9997 | 0.9754 | 5 |  | X | X | X | NO |
| Hetgly.G000014464 | Hetgly02651 | 0.9997 | 0.9599 | 5 | X |  |  |  | NO |
| Hetgly.G000014527 | Hetgly09638 | 0.8601 | 0.8579 | 2 | X |  |  |  | NO |
| Hetgly.G000015939 | Hetgly07801 | 0.9998 | 0.9350 | 1 |  |  |  | X | NO |
| Hetgly.G000016234 | Hetgly16974 | 0.9808 | 0.9515 | 7 |  |  |  | X | NO |
| Hetgly.G000016328 | Hetgly13997 | 0.9994 | 0.9480 | 4 |  |  | X | X | YES<sup>1</sup> |
| Hetgly.G000016675 | Hetgly05522 | 0.9997 | 0.9099 | 3 | X |  |  |  | NO |
| Hetgly.G000016740 | Hetgly11688 | 0.9997 | 0.9764 | 2 |  | X |  | X | NO |
| Hetgly.G000016899 | Hetgly14389 | 0.8984 | 0.6139 | 6 | X |  |  |  | NO |
| Hetgly.G000017118 | Hetgly11576 | 0.9998 | 0.9692 | 2 | X |  |  |  | YES<sup>1</sup> |
| Hetgly.G000017651 | Hetgly09041 | 0.9963 | 0.8390 | 2 |  |  |  | X | NO |
| Hetgly.G000017808 | Hetgly20463 | 0.9998 | 0.8331 | 9 |  |  | X | X | NO |
| Hetgly.G000018759 | Hetgly06909 | 0.9997 | 0.9492 | 1 |  |  |  | X | NO |
| Hetgly.G000018760 | Hetgly06908 | 0.9997 | 0.9683 | 1 |  | X |  |  | YES<sup>1</sup> |
| Hetgly.G000018896 | Hetgly03920 | 0.9998 | 0.9405 | 3 |  | X |  |  | NO |
| Hetgly.G000019218 | Hetgly20700 | 0.9998 | 0.9576 | 8 |  |  | X | X | NO |
| Hetgly.G000021894 | Hetgly08087 | 0.9985 | 0.7620 | 1 | X |  |  |  | YES<sup>1</sup> |
| Hetgly.G000022800 | Hetgly08816 | 0.9998 | 0.9282 | 1 |  | X | X | X | YES<sup>1</sup> |
| Hetgly.G000023464 | Hetgly12039 | 0.9998 | 0.8796 | 4 | X |  |  |  | NO |
| Hetgly.G000028400 | Hetgly03197 | 0.9997 | 0.9618 | 3 |  | X | X | X | NO |

## Discussion

The main objective of this study was to investigate the defense mechanism of PI 437654, a highly resistant soybean line (FI=0%) against SCN HG type 1.2.5.7. PI 548402 and PI 88788, which exhibit lower levels of resistance (FI=10% and 63%, respectively), were also included to validate resistant genes and identify the resistance genes unique to PI 437654. Furthermore, the roots of these three resistant lines, as well as the susceptible line Lee 74 (FI=100%), were analyzed to study changes in soybean root transcriptome in response to SCN. To identify candidate genes responsible for resistance in PI 437654, GO enrichment analyses were employed, which allowed for the extraction of gene sets involved in the enriched biological process. These findings provide insight into the mechanisms underlying resistance to SCN, particularly in PI 437654, and can aid in the development of more resistant soybean cultivars. Simultaneously, dual RNA sequencing allows us to examine SCN reactions across multiple soybean hosts (e.g. compatible, semi-compatible, semi-incompatible, incompatible) to study the expression of SCN pathogenicity genes during the infection process.

In this research a long-term SCN stress at 5 and 10 dpi was chosen for RNA sequencing analysis to investigate the dual transcriptome reaction between soybean and SCN. Hammond et al. (2004) classified genes that respond to stress into two categories: “early” and “late” genes. The “early” genes exhibit a rapid response and are generally non-specific to target stresses, while the “late” genes have a delayed expression but can have a significant impact on the morphology, physiology, and/or metabolism of plants. Furthermore, these “late” genes are often specific to target stresses. According to Matsye et al. (2011), Peking-type resistance is characterized by a rapid and potent resistant reaction that results in the formation of a necrotic region around the syncytium by 5 dpi. On the other hand, PI 88788-type resistance is a prolonged but potent resistant reaction, which does not show any cytological evidence of a reaction at 5 dpi. Previous studies on cell fate against SCN have found that selecting “late” genes increases the likelihood of identifying specific genes. Consequently, 5 and 10 dpi was used for dual transcriptome reaction between soybean and SCN. The results of our dual transcriptome analysis provide valuable insight into the complex reaction of soybean gene expression in response to SCN parasitism. Furthermore, the coordinated expression of SCN HG type 1.2.5.7 genes potentially involved in parasitism against different soybean lines can now be better understood.

According to the total number of reads mapped to *H. glycine*, results indicated that SCN HG type 1.2.5.7 had the lowest penetration in PI 437654, possibly due to a strong passive defense mechanism, while it had the highest penetration in Lee74, likely due to a physical barrier weakness. A significant decrease in the frequency of reads in Lee74 at 10 dpi suggested that SCN HG type 1.2.5.7 had overcome the defense mechanism of Lee74, leading to successful development into adult females and males. In PI 548402, the observed reduction in read frequency may be attributed to the strong two-layered immunity system, which overcomes SCN HG type 1.2.5.7. It is possible that the second-stage juveniles (J2) nematode cannot form specialized feeding sites, syncytia, and, therefore, are unable to copulate, leading to their eventual starvation and death. At 10 dpi, while the total number of reads mapped to *H. glycine* in PI 88788 was significantly reduced, it was still remained relatively large. This observation may suggest that there was a battle between the SCN and PI 88788, where neither was able to fully overcome the other.

Comapring differentially expressed genes (DEGs) within each genotype under infested and non-infested conditions at two time points, 5 & 10 dpi, demonstrated the number of DEGs significantly decreased in PI 437654, PI 548402, and PI 88788 at 10 dpi. In contrast, the number of DEGs remarkably increase in Lee74, which may be due to the ability of SCN effector proteins to hijack and alter the gene expression patterns of Lee74, leading to the induction of a large number of DEGs.

Host transcriptional pattern reprogramming is often triggered by pathogen invasion. Plasma membrane Ca channels are among the early sensors that respond to pathogen attacks by increasing Ca influx into the cell cytoplasm, as reported by Sun et al. (2015). Ca2+ sensor proteins, such as calmodulin, EF-hand domain, and Ca2+-dependent protein kinases (CDPKs), detect the transient increase in Ca2+ signatures, as observed by Houqing Zeng et al. (2017) and Sun et al. (2015). Calmodulin, despite lacking enzymatic activity, binds to calmodulin-binding proteins, thereby stimulating the synthesis and accumulation of SA during immunity, as highlighted by Choudhury et al. (2017) and Gilroy et al. (2016). SA, in turn, activates systemic acquired resistance (SAR), providing broad-spectrum and long-lasting resistance against pathogens (Sun et al., 2015). The study identified calmodulin-binding proteins *(Glyma.19G229500, Glyma.03G232400,* and *Glyma.19G229400)* at 5 dpi and calmodulin-binding proteins (*Glyma.19G229500, Glyma.05G237200, Glyma.07G093900,* and *Glyma.09G182400*) at 10 dpi in PI 437654, but not in Lee74. *Glyma.03G232400* and *Glyma.19G229400* were unique to PI 437654, and *Glyma.05G237200* and *Glyma.07G093900* were also observed in PI 88788 (Figure 7). *Glyma.19G229500* and *Glyma.09G182400* were validated in both PI 548402 and PI 88788, with *Glyma.19G229500* being differentially expressed in both time points and validated by other resistant lines. These findings align with previous studies by Kofsky et al. (2021) and Zhang et al. (2017) that reported the presence of calmodulin binding proteins in transcriptome comparisons of different genotypes under SCN HG type 2.5.7-treated conditions. Additionally, the study detected two differentially expressed EF-hand domain proteins (*Glyma.19G160100* and *Glyma.16G214800*) in PI 437654 at 5 dpi, but not in other resistant lines (Table 2 & Figure 7).

**Figure 7.**
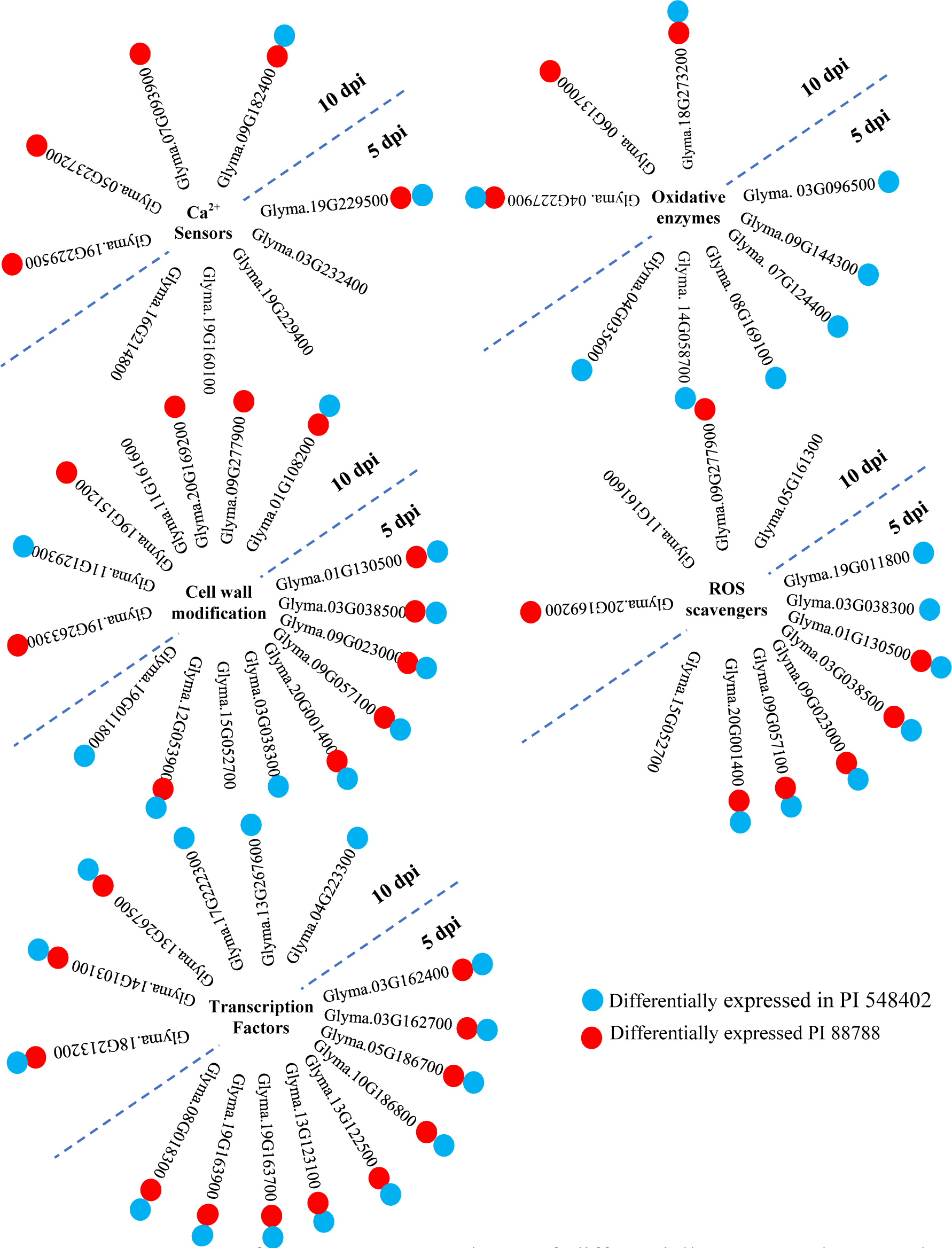
A summary of intergenotype analyses of differentially expressed genes that were significantly upregulated in PI 437654 in response to SCN HG type 1.2.5.7 at 5 and 10 dpi and were not observed in Lee74 as a susceptible line. Each cluster of genes is associated with specific functional categories involved in SCN-resistant defense mechanisms. Blue balls indicate SCN-responsive genes also expressed in PI 548402. Red balls indicate SCN-responsive genes also expressed in PI 88788.

Protein Kinase is a critical component of intracellular signal transduction and plays an essential role in stress response (Wang et al., 2015). In current research, three proteins with protein kinase activity, namely *Glyma.07G184000, Glyma.17G173000,* and *Glyma.18G219600*, were identified at 5 dpi. Among them, *Glyma.07G184000* and *Glyma.17G173000* were validated using both PI 548402 and PI 88788, while *Glyma.18G219600* was only observed as differentially expressed in PI 437654. These protein kinases identified as SCN-responsive genes in the current study are aligned with the observations made by Ithal et al., 2007, and Han et al., 2015, which also detected these genes in response to SCN HG type 0 and HG type 1.2.3.5.7 (Ithal et al., 2007; Han et al., 2015)

Transcription factors play crucial roles in signal transduction by activating or suppressing the expression of defense genes and regulating the crosstalk between different signaling pathways. As they bind to specific cis-acting elements in gene promoters, transcription factors are positioned at the penultimate step of the signal cascade to directly control the downstream target gene expression. In the current study, several proteins with AP2/ERF domains, such as *Glyma.03G162400, Glyma.03G162700, Glyma.05G186700, Glyma.10G186800, Glyma.13G122500, Glyma.13G123100, Glyma.19G163700,* and *Glyma.19G163900*, which have transcription regulator activity, were found to be differentially upregulated in PI 437654 at 5 dpi. These genes were confirmed to play an important role against SCN HG type 1.2.5.7 through their expression in other resistant lines, including PI 548402 and PI 88788. The presence of AP2 is aligned with the observations made by Kofsky et al., who studied resistant and susceptible resources from *G. soja* against SCN HG type 2.5.7. Another upregulated transcription factor in all three resistant lines at 5 dpi is the WRKY protein (e.g., *Glyma.08G018300*). At 10 dpi, six WRKY genes, including *Glyma.04G223300, Glyma.13G267600, Glyma.17G222300, Glyma.13G267500, Glyma.14G103100,* and *Glyma.18G213200*, were detected in PI 437654. Of these, the first three were also observed in PI 88788, and the last three were validated in both PI 548402 and PI 88788. The upregulation of WRKY genes as SCN-responsive genes in this study is supported by Zhang et al., 2017, as well as the observations made by Ithal et al., 2007, Han et al., 2015, and Kofsky et al., 2021.

Plants encounter various stresses and ROS such as hydrogen peroxide (H_2_O_2_), superoxide anions (O_2_•−), hydroxyl radical (OH•), and singlet oxygen (1O_2_) play important roles in signal transduction. However, excessive ROS accumulation can be harmful and even lead to cell death. Thus, a delicate balance between ROS production and ROS-scavenging pathways must be maintained (Jun You and Zhulong Chan, 2015). In the present study, at 5 dpi, several peroxidase genes including *Glyma.19G011800, Glyma.01G130500, Glyma.03G038300, Glyma.03G038500, Glyma.09G023000, Glyma.09G057100, Glyma.15G052700,* and *Glyma.20G001400* were found to be differentially expressed in PI 437654, but not in Lee74. Among these peroxidase genes, *Glyma.01G130500, Glyma.03G038500, Glyma.09G023000, Glyma.09G057100,* and *Glyma.20G001400* were observed in both PI 548402 and PI 88788, while *Glyma.19G011800* and *Glyma.03G038300* were only validated in PI 548402. *Glyma.15G052700* was the only unique peroxidase gene that was differentially expressed in PI 437654 at 5 dpi. At 10 dpi, *Glyma.09G277900, Glyma.20G169200,* and *Glyma.11G161600* were identified as peroxidase genes, while *Glyma.05G161300* was differentially expressed and annotated as a Glutathione S-transferase (GST). The first two peroxidase genes were confirmed using PI 88788, while the last peroxidase and GST were only identified in PI 437654. These findings are in agreement with the observation by Miraeiz et al., 2020, who studied RNA-Seq profiling of Peking, Fayette, Williams 82, and a wild relative (*Glycine soja*, PI 468916) against SCN HG type 0.

Oxidative enzymes are pivotal to plant metabolism, catalyzing a wide array of reactions involved in hydroxylation, DNA repair, post-translational modification, as well as the activation and catabolism of plant growth regulators. The 2OG-Fe(II) oxygenase superfamily and Cytochrome P450 (P450) are two important classes of these enzymes (Farrow and Facchini, 2014). In this investigation, four genes belonging to the 2OG-Fe(II) oxygenase superfamily (*Glyma.03G096500, Glyma.07G124400, Glyma.08G169100,* and *Glyma.14G058700*) were identified at 5 dpi, and two genes (*Glyma.04G227900* and *Glyma.18G273200*) were detected at 10 dpi in PI 437654, but not in Lee74 (Figure 7). Additionally, 11 genes annotated as Cytochromes P450 were identified at 10 dpi in PI 437654 (e.g., *Glyma.01G135200, Glyma.02G156100, Glyma.03G143700, Glyma.05G022100, Glyma.09G049200, Glyma.10G114600, Glyma.11G062500, Glyma.11G062600, Glyma.11G062700, Glyma.15G156100,* and *Glyma.16G195600*) (Table 3). This study’s results reveal that only two P450 genes (*Glyma.04G035600* and *Glyma.09G144300*) were validated and differentially expressed in PI 548402 but not in Lee74 (Figure 7 and Table 4), implying that these genes play a crucial role in soybean defense against SCN infection. Our findings are in agreement with previous studies by Ithal et al., 2007, and Han et al., 2015, further highlighting the importance of oxidative enzymes in the plant’s defense response.

The plant cell wall can serves as a protective and physical barrier for limiting pathogen penetration into the plant cell (McCann and Roberts, 1991; Carpita and Gibeaut, 1993). Lignin, a three-dimensional polymer, is a major component of plant cell walls, composed of monomeric units such as syringyl, guaiacyl, and p-hydroxyphenyl, derived from phenylalanine in the cytoplasm and transported to the cell wall. The monomeric units are polymerized by laccase and peroxidase enzymes, which catalyze the random cross-linking necessary for the formation of the lignin polymer (Vanholme et al., 2010; Tronchet et al., 2010; Hamann, 2012). Polymerized lignin reinforces the strength and rigidity of plant cell walls and is a key component of the plant’s response to environmental factors (Lee et al., 2019; Billa et al., 1997). In this study, nine differentially expressed peroxidase genes were overrepresented in PI 437654 at 5 dpi (Table 2). Based on KEGG analyses, these genes are involved in the phenylpropanoid pathway and are directly responsible for producing syringyl lignin, guaiacyl lignin, 5-hydroxy-guaiacyl lignin, and p-hydroxyphenyl lignin. Among these genes, *Glyma.01G130500*, *Glyma.03G038500*, *Glyma.09G023000*, *Glyma.09G057100*, and *Glyma.20G001400* were validated through PI 548402 and PI 88788 and were not found to be differentially expressed in Lee 74. *Glyma.03G038300* and *Glyma.19G011800* were only validated in PI 548402 and were not observed in the susceptible line (Figure 7 & Table 4). Another peroxidase gene that was differentially expressed at 5 dpi was *Glyma.15G052700*, which was only observed in PI 437654 and not in the susceptible line (Figure 7 & Table 4). At 10 dpi, one differentially expressed Laccase gene and nine peroxidase genes were detected. *Glyma.01G108200* was annotated as a laccase gene and was validated by both PI 548402 and PI 88788. Out of the nine peroxidase genes, only *Glyma.09G277900* and *Glyma.20G169200* were not found to be differentially expressed in Lee74 and were validated only by PI 88788 (Figure 7 & Table 5). Another peroxidase gene that was differentially expressed at 10 dpi was *Glyma.11G161600*, which was only observed in PI 437654 and did not significantly upregulate in Lee74 (Figure 7 & Table 3). In addition, Dirigent is another cell-wall-related gene that modulates cell-wall metabolism (Li N. et al., 2017). At 10 dpi, Glyma.19G151200, which was annotated as a dirigent gene, was not detected in the susceptible line and was validated in PI 88788 (Figure 7). Our finding about the role of peroxidase, laccase, and dirigent in cell wall rigidity against SCN invasion is supported by Afzal et al., 2009, which compared two NILs including rhg1rhg1/Rhg4Rhg4 and Rhg1Rhg1/Rhg4Rhg4 against SCN HG type 0 and Miraeiz et al., 2020 who studied different soybean lines against SCN HG type 0. Glycoside hydrolase is another gene involved in cell wall polysaccharide metabolism (Minic, 2008). Among the DEGs observed during SCN invasion in PI 437654, *Glyma.12G053900* at 5 dpi and *Glyma.11G129300*, *Glyma.12G054200*, and *Glyma.13G346700* at 10 dpi were upregulated. All of these genes are annotated as glycoside hydrolases. *Glyma.12G053900* was observed in both PI 548402 and PI 88788, and *Glyma.11G129300* was validated through PI 548402. Lipoxygenase is another gene that induces cell wall modification to limit pathogen invasion (Vellosillo et al., 2013), and at 10 dpi, the lipoxygenase gene *Glyma.19G263300* was detected in PI 437654 and validated in PI 88788.

Interestingly, the rhg1 genes (e.g., *Glyma18g02580, Glyma18g02590,* and *Glyma18g02610*) and Rhg4 gene (*Glyma.08G108900*), which are promising genes known to confer SCN resistance to HG type 0, were not identified as significantly different in the current study, consistent with the findings of Zhang et al., 2017. Additionally, the pathways of plant-pathogen interaction, carbon fixation in the photosynthetic organism, and carbon metabolism, which were the top three enriched pathways for up-regulated DEGs in PI 437654, were not observed in other resistant lines. This finding is consistent with the observation by Shi et al., 2021, who studied PI 437654 against SCN HG type 1.2.3.5.7. However, this result is in contrast to the reports of Zhang et al., 2017, who studied on *Glycine soja* interaction with SCN HG type 2..5.7 and identified genes involved in carbon fixation and photosynthesis pathways as remarkably downregulated. According to Shi et al., 2021, a large number of DEGs that were upregulated in the incompatible soybean variety PI 437654 were involved in the plant hormone pathway, MAPK signaling, and phenylpropanoid biosynthesis pathway, which is in agreement with the results of the current study on PI 437654.

Dual RNA sequencing has provided a significant advantage in investigating the interaction between pathogen and host simultaneously. Through transcriptome analysis of SCN HG type 1.2.5.7, novel secreted effectors with potential roles in the SCN-host interaction have been identified. The establishment and maintenance of the syncytium, a crucial step for the long-term parasitic success of SCN, involves multiple stages such as hatching stimuli, host attraction, root penetration, tissue modification, feeding site formation, and immune system suppression. The pathogenicity and severity of SCN are dependent on the successful completion of these stages, as well as the pathogen’s classification as pathogenic or non-pathogenic. This study represents the first investigation of SCN transcriptome in different hosts with varying levels of resistance, shedding light on the virulence strategies employed by SCN to overcome resistant hosts. Notably, 51 putative effectors showing differential expression patterns in both resistant and susceptible lines were identified, with 39 of them being newly discovered. Furthermore, only a limited overlap was observed between the effectors identified in PI 437654 and those identified in other resistant and susceptible lines, highlighting the adaptive ability of SCN to modulate its virulence genes to overcome resistant hosts.

## Conclusion

Our study has uncovered the mechanisms underlying the robust resistance of PI 437654 against SCN HG type 1.2.5.7, which involves reinforcement of cell walls and enhancement of physical barriers that impede pathogen penetration. Notably, PI 437654 exhibits a lower number of defense-related genes, suggesting its ability to confront H. glycine at the root entry point. Our analysis of egg production in different hosts against SCN HG type 1.2.5.7 has led to the classification of PI 437654 as an incompatible line, PI 548402 as a semi-incompatible line, PI 88788 as a semi-compatible line, and Lee74 as a compatible line, providing a spectrum of resistant lines for investigating the soybean-SCN interaction. By focusing on PI 437654 and validating with PI 548402 and PI 88788, we have identified several promising candidate genes for SCN resistance, including 2OG-Fe(II) oxygenase superfamily, cytochrome P450, AP2 domain, WRKY, protein kinase, peroxidase, leucine-rich repeat, calmodulin-binding protein, laccase, and dirigent, consistent with previous studies. As SCN resistance is a complex trait, it is likely that multiple genes are required to confer resistance against *H. glycines*. Furthermore, our findings support the hypothesis that the SCN genome contains multiple hot spot regions for non-synonymous mutations, driving its adaptive ability to overcome resistance in different hosts. The diverse behavior of SCN HG type 1.2.5.7 in different hosts, as observed through transcriptome monitoring validates this hypothesis. It is plausible that secreted proteins or effectors of SCN undergo accelerated evolution in response to strong selective pressure from host immunity. These findings may explain the shift in the SCN population in soybean fields where the same source of resistance has been repeatedly used.

Our study has uncovered new molecular mechanisms that underlie soybean’s resistance to SCN, and has deepened our understanding of the complex interplay between SCN and soybean hosts. These discoveries have far-reaching implications for the development of effective strategies to manage SCN and underscore the importance of breeding for resistance by targeting multiple genes. Future research in this area holds promise for identifying additional mechanisms and genes that contribute to SCN resistance, ultimately improving soybean production and sustainability in regions affected by SCN.

## Methods

### Soybean and SCN procurement for inoculation

The SCN HG type 1.2.5.7 inbred strain provided by the University of Illinois was maintained for more than 50 generations. To extract eggs, cysts were first sterilized by briefly treated with 0.5% sodium hypochlorite for approximately 5 seconds, followed by thorough rinsing with tap water. Eggs were extracted by crushing cysts with a rubber stopper and collecting them using a nested sieve consisting of 75 µm (mesh 200) and 25 µm (mesh 500) sieves (USA Standard Testing Sieve), following which the number of eggs per unit volume was determined by stirring the egg suspension with a magnetic stir bar and counting an aliquot of the suspension using a 10x microscope on a chambered counting slide. This process was repeated eight times to estimate the number of eggs per unit volume.

Seeds of PI 88788, PI 437654, PI 548402, and Lee-74 were procured from Agriculture and Agri-Food Canada, Harrow, Canada. The soybean seeds were surface-sterilized by soaking in a 5% (v/v) sodium hypochlorite for 5 minutes and washed with sterilized water three times for 2 minutes each time. The sterilized seeds were then germinated between moist sterilized filter papers in a petri dish at the dark condition and 25 °C. After three days, the seedlings were transferred to prepared cons that contained a cotton ball, turface, and a mix of two types of sand (A 50:50 mix of masonry sand and beach sand) from bottom to top. One day after the transfer, each plant was inoculated using 4000 eggs of SCN HG type 1.2.5.7 and incubated in a growth chamber at the University of Guelph. The chamber was set to a day/night temperature was set to 27 °C/25 °C with a photoperiod of 16 hours and light intensity of 450 umoles/m2/s, and the relative humidity was set at 60%. The seedlings were watered daily to maintain plant moisture. This experiment followed a complete block design (CRD) with three biological replications.

### Female Index (FI) calculation

At 52 dpi, the cysts were thoroughly washed with water through an 850 µm (mesh 20) sieve stacked on a 250 µm (mesh 60) sieve (USA Standard Testing Sieve). The Female index (FI) was calculated on a per-plant basis using the following equation:

Female Index (FI) = (Number of cysts on test line)/(Number of cysts on Lee74) X 100

Following the cyst count and FI% calculation, eggs were released from the cysts and counted under a 10x microscope (Olympus Canada Inc).

### *rhg1* and *Rhg4* copy number variation

To determine the copy number of *rhg1* and *Rhg4*, the GmSNAP18 and GmSHMT08 genes were used as references, following the protocols described by Kadam et al., 2016 and Patil et al., 2019. Oligonucleotide probes specific for *rhg1* and *Rhg4* were labelled with FAM at the 5′ ends, while the probe for the reference gene, lectin (*Le1*), was labelled with the fluorescent dye VIC at the 5′ ends. The 3′ ends of all probes were labelled with the quencher dye MGBNFQ (Table S12; Applied Biosystems, Foster City, Calif).

Real-time PCR was performed using a 10 μl reaction mixture containing 5 μl 2x TaqMan Universal PCR Master Mix (Applied Biosystems), 500 nM of each primer, 100 nM of each probe, and 15ng of genomic DNA. Two technical replicates were conducted for each sample, and template-free or negative controls were included. The QuantStudio 6 flex System (Applied Biosystems, CA) was used to run the real-time PCR program, consisting of an initial denaturation step of 10 min at 95 °C, 40 cycles of 15 s at 95 °C, and 1 min at 60 °C. The copy number assay was calibrated using Williams 82 as a control sample, which carries a single-copy of insertion. Copy number analysis of the target genes was performed using the CopyCaller Software v2.0 (Applied Biosystems), following the manufacturer’s instructions.

### Tissue collecting and Infection Confirmation

The current study aimed to identify specific SCN-responsive genes in soybean and host-specific pathogenesis genes in *H. glycine*, and 5 and 10 dpi were chosen as the optimal time points based on a previous study (Matsye et al. 2011; Hammond et al., 2004). Root samples were collected from both inoculated and control seedlings, rapidly washed, and flash-frozen quickly using liquid nitrogen. The samples were then stored at -80 °C for subsequent analysis. To confirm the infection of the roots with SCN HG type 1.2.5.7 at 5 and 10 dpi, PCR amplification of the nematode 18S ribosomal gene was performed using root-extracted DNA (Table S13).

### RNA extraction and Dual RNA sequencing

Total RNA was extracted from both soybean and SCN using the Invitrogen™ PureLink™ RNA Mini Kit, following the manufacturer’s instructions. The extract RNA samples were evaluated for both quantity and quality of at the Genomics Facility of the University of Guelph (Guelph, ON) using an Agilent 2100 BioAnalyser System (Agilent Technologies, Palo Alto, CA). Only high-quality RNA samples (RNA Integrity Number ≥ 9) were selected for sequencing at the Genome Quebec, Innovation Center of McGill University (Montreal, QC).

The raw data were subjected to quality control checks, which involved removing reads containing adaptor sequences, poly-N sequences, and low-quality reads. Reference genome and gene model annotation files were obtained from the Phytozome website (Goodstein, D. M. et al., 2012) for soybean and WormBase Parasite website (Howe, K. L. et al., 2016 & 2017) for SCN.

The reads were aligned to reference genome using Hisat2 v. 2.0.5 (Liao et al., 2014), and FeatureCounts v. 1.5.0-p3 (Mortazavi et al., 2008) was utilized to quantify the number of reads mapped to each gene. Gene expression levels were measured in FPKM values. Differentially expressed genes were identified using DEBRowser/DESeq2 with log2 FC (fold change) of >1 and a p-value of < 0.05.

### Data analyses and gene pathway analyses

The jvenn software (Bardou et al. 2014) was employed to generate Venn diagrams. For gene annotation analysis of soybean and SCN, the Biomart in EnsemblPlants database (accessible at https://plants.ensembl.org/) and Wormbase Parasite were used, respectively. GOE analysis, hierarchical clustering tree, network analysis, and KEGG were performed using ShinyGO (ShinyGO v0.61) (Ge et al. 2020).

### Putative Effectors

To identify putative effectors among the differentially expressed genes in SCN, we evaluated the presence of secretory signal peptide and the probability of extracellular localization for all differentially expressed proteins. First, we extracted the protein sequences of each differentially expressed gene from Wormbase Parasites (Howe, K. L. et al., 2016 & 2017). Then, we used SignalP 6.0 (https://services.healthtech.dtu.dk/service.php?SignalP) to predict signal peptides and cleavage sites, with default Eukaryote setting, based on the protein language model (Teufel et al., 2022). To further filter the candidate effectors, we utilized the DeepLoc 2.0 server (https://services.healthtech.dtu.dk/service.php?DeepLoc-2.0) (Vineet Thumuluri et al., 2022) to predict extracellular localization. Proteins showing at least a 50% probability of containing a signal peptide signature and extracellular localization prediction were selected and considered as putative effectors.

## Acknowledgments

The Authors would like to acknowledge the technical assistant of Cheryl Van Herk, at Ontario Ministry of Agriculture, Food and Rural Affairs, Ridgetwon, ON, Canada, and also Derek Lawrence, at Harrow Research and Development Centre, Agriculture and Agri-Food Canada, London, ON, Canada. The Authors also thanks Dr. Alison Colgrove from Department of Crop Sciences at University of Illinois, US for provding us with SCN HG type 1.2.5.7.

## Author contributions

M.E., conceived, conceptualized, planned, and suppervised the study. S.T. contriubted in conceptualizing and planning the study, performed the experiments, collected the samples, and conducted most of the analyses. S.S., and S.T., performed RNA-seq analyses and the vitualization. A.T. contributed in SCN bioassay. O.W. provided IP germplasm. J.G.M, A.T., O.W., and D.T., contributed in methodology and experiment design. All authors read and approved the final manuscript.

## Supplemental data

The following materials are available in the online version of this article.

**Supplemental Figure S1.** Amplifying the SCN 18S ribosomal gene from DNA extracted from the roots.

**Supplemental Table S1** Number of reads mapped to *Glycine ma*x and *Heterodera Glycines*

**Supplemental Table S2** GO biological analyses of upregulated genes in PI 437654 at 5 dpi

**Supplemental Table S3** GO biological analyses of upregulated genes in PI 437654 at 10 dpi

**Supplemental Table S4** Go term categories of genes that uniquely upregulated in PI 437654 at 5dpi

**Supplemental Table S5** GO term categories of genes that identified in PI 437654 and PI 548402 at 5 dpi

**Supplemental Table S6** GO enrichment analyses of SCN-responsive genes overlapped in PI 437654, PI 548402 and PI 88788 at 5 dpi

**Supplemental Table S7** GO enrichment analysis unique DEGs in PI 437654 at 10 dpi

**Supplemental Table S8** Go enrichment analysis of DEGs that overlapped in PI 437654 and PI 548402 at 10 dpi

**Supplemental Table S9** GO enrichment analysis of SCN-responsive genes that overlapped in PI 437654 and PI 88788 at 10 dpi

**Supplemental Table S10** The pathways analyses of DEGs in PI 437654, PI 548402, and PI 88788 at 5dpi

**Supplemental Table S11** The pathways analyses of DEGs in PI 437654, PI 548402, PI 88788 and Lee74 at 10 dpi

**Supplemental Table S12** Sets of primers and probes used for TaqMan assay to evaluate copy number variation (CNV) of *rhg1* and *Rhg4* genes

**Supplemental Table S13** Primer sequences of nematode 18S ribosomal gene

---

Figure S1- Amplifying the SCN 18S ribosomal gene from DNA extracted from the roots

**Figure 1S.**
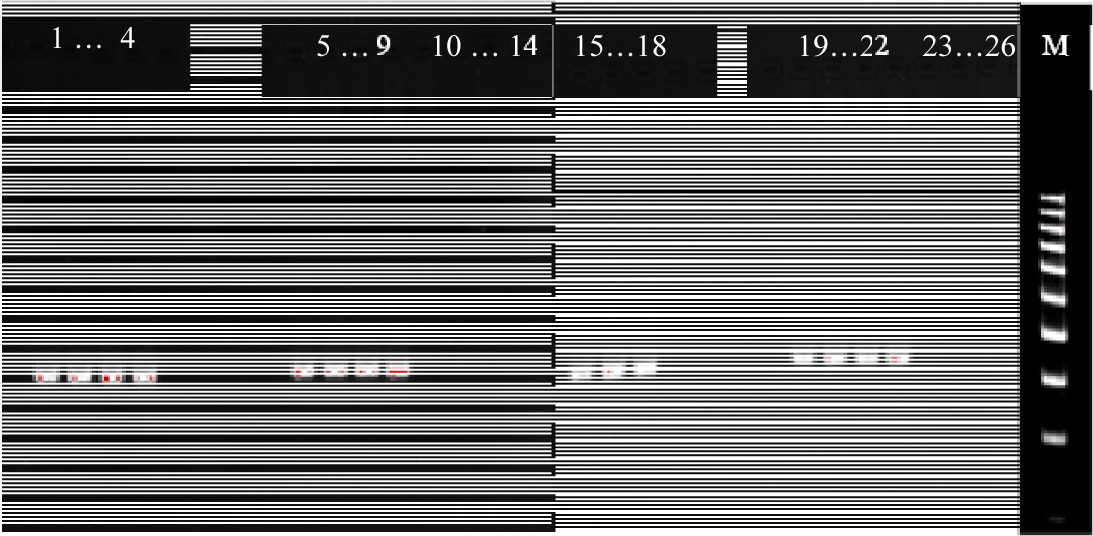
PCR amplification of 18S ribosomal RNA (250 bp band) in SCN, infected plants and Control at 5 and 10 dpi. Lanes 1 to 4 and Lane 15 to 18 are SCN. Lanes 5 to 9 are infected plants at 5 dpi. Lane 10-14 are Control plants at 5 dpi. Lane 19 to 22 are infected plants at 10 dpi. Lane 23 to 26 are control plants at 10 dpi.

